# Lipid Oxidation Controls Peptide Self-Assembly near Membranes

**DOI:** 10.1101/2022.08.02.502408

**Authors:** Torsten John, Stefania Piantavigna, Tiara J. A. Dealey, Bernd Abel, Herre Jelger Risselada, Lisandra L. Martin

## Abstract

The self-assembly of peptides into supramolecular fibril structures has been linked to neurodegenerative diseases such as Alzheimer’s disease but has also been observed in functional roles. Peptides are physiologically exposed to crowded environments of biomacromolecules, and particularly membrane lipids, within a cellular milieu. Previous research has shown that membranes can both accelerate and inhibit peptide self-assembly. Here, we studied the impact of biomimetic membranes that mimic cellular oxidative stress and compared this to mammalian and bacterial membranes. Using molecular dynamics simulations and experiments, we propose a model that explains how changes in peptide-membrane binding, electrostatics, and peptide secondary structure stabilization determine the nature of peptide self-assembly. We explored the influence of zwitterionic (POPC), anionic (POPG) and oxidized (PazePC) phospholipids, as well as cholesterol, and mixtures thereof, on the self-assembly kinetics of the amyloid β (1–40) peptide (Aβ_40_), linked to Alzheimer’s disease, and the amyloid-forming antimicrobial peptide uperin 3.5 (U3.5). We show that the presence of an oxidized lipid had similar effects on peptide self-assembly as the bacterial mimetic membrane. While Aβ_40_ fibril formation was accelerated, U3.5 aggregation was inhibited by the same lipids at the same peptide-to-lipid ratio. We attribute these findings and peptide-specific effects to differences in peptide-membrane adsorption with U3.5 being more strongly bound to the membrane surface and stabilized in an α-helical conformation compared to Aβ_40_. Different peptide-to-lipid ratios resulted in different effects. Molecular dynamics simulations provided detailed mechanistic insights into the peptide-lipid interactions and secondary structure stability. We found that electrostatic interactions are a primary driving force for peptide-membrane interaction, enabling us to propose a model for predictions how cellular changes might impact peptide self-assembly *in vivo*, and potentially impact related diseases.

## Introduction

The self-assembly of peptides in a physiological environment into supramolecular fibril structures has been implicated in ageing-related and neurodegenerative diseases.^1^ One example is amyloid β peptide (Aβ) that aggregates in the brains of patients diagnosed with Alzheimer’s disease.^2,3^ However, peptide fibrils have not only been related to disease but were identified as functional, non-pathological states, and have developed structural advantages as functional materials, such as found in spider silk.^4,5^ The fibril-forming peptide uperin 3.5 (U3.5) was first isolated as an antimicrobial peptide (AMP) and may be related to the innate immune system of the Australian toadlet *Uperoleia mjobergii*.^6–8^ Peptide fibrils are typically water-insoluble and form a common cross-β sheet structure, as observed by electron microscopy and X-ray diffraction.^9,10^ Recent cryoEM studies also identified cross-α fibril structures for a number of peptides.^11–14^

The formation of peptide fibrils follows typical nucleation-elongation kinetics with a slow nucleation phase followed by a fast elongation and growth of the peptide oligomers into fibrillar aggregates (see Figure 1a).^15,16^ Several studies suggested an α-helical peptide conformation as an intermediate toward β-sheet rich fibrils.^17–20^ However, the physiological role of amyloid peptides and the biochemical processes that cause aggregation and disease are still under investigation.^21,22^ Since antimicrobial properties have not only been found for U3.5 but also for the Alzheimer-related Aβ peptide,^23^ studies suggested links between the antimicrobial activity of peptides and their aggregation into amyloid fibrils.^8,23–25^

**Figure 1.**
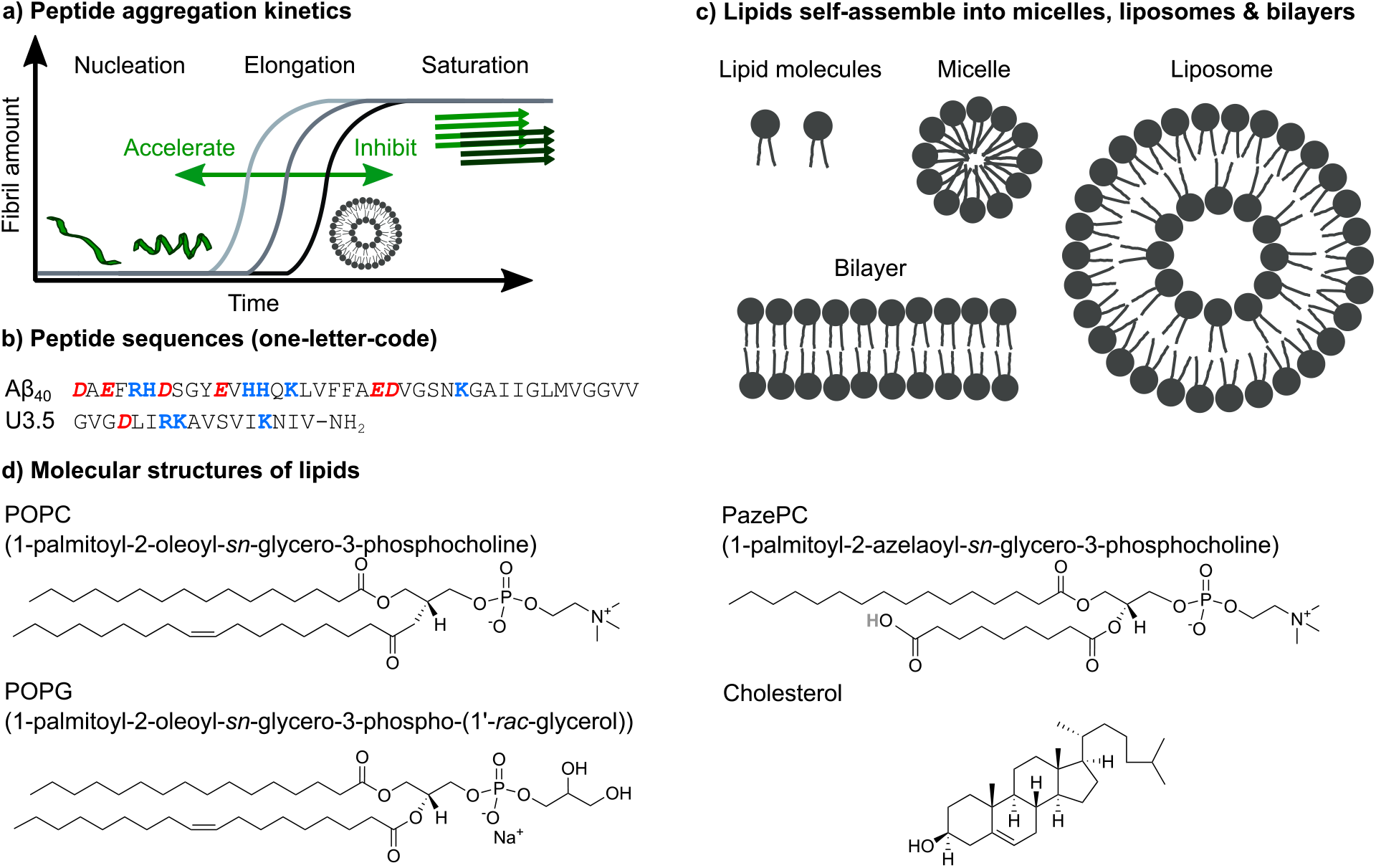
Overview of the impact of lipids on peptide self-assembly into fibrils. (a) Typical nucleation-elongation kinetics of fibril formation with the slow formation of critical nuclei and subsequent rapid fibril growth. The presence of lipids accelerates or inhibits peptide fibril formation, resulting in shorter or longer times for the nucleation phase, respectively. (b) Peptide sequences of Aβ_40_ and U3.5 in one-letter code (acidic groups: red and bold italics, basic groups: blue and bold). (c) Phospholipids with hydrophilic head groups and hydrophobic tails typically spontaneously self-assemble into lipid bilayers, micelles or liposomes. (d) The chemical structures of the studied phospholipids POPC, POPG and PazePC as well as of cholesterol are shown. The carboxyl group of PazePC may be (partially) deprotonated under experimental condition.

Under physiological conditions, peptides are surrounded by other biomacromolecules in crowded environments and cell membranes play an important role.^26–29^ This is particularly relevant as the pathology of amyloidogenic peptides has been linked to their peptide membrane activity.^30–32^ Both the impact of membranes on peptide structure and self-assembly kinetics as well as the action of peptides on membranes have been studied extensively.^33,34,43,35–42^ The membrane damage caused by amyloidogenic peptides has been attributed to different oligomeric species as well as the fibril growth process.^44,45^ Previous work either proposed the disruption of membranes by peptides or the modulation of peptide self-assembly by membranes as the initial process in relation to disease, or considered both processes as concomitant.^33,35,36,46,47^ Sparr and Linse emphasized the important role of membrane properties, and particularly the protein-to-lipid ratio, among the factors contributing to lipid-protein interactions in amyloid formation.^48^

In this work, we focused our attention on the role of the oxidized membrane lipid PazePC on the structure and self-assembly kinetics of the amyloid β (1–40) peptide (Aβ_40_)^2,3,49^ and the antimicrobial peptide uperin 3.5 (U3.5) (Figure 1b), aggregating near model membranes.^6–8^ While Aβ_40_ is a widely studied peptide related to Alzheimer’s disease,^2^ U3.5 has originally been identified as an AMP. AMPs are generally cationic and known to adopt an α-helical conformation when in contact with membranes surfaces, stabilizing either intermediates toward peptide fibrils or off-pathway oligomers.^17,50–53^ Interestingly, the membrane disruption activity of antimicrobial and amyloidogenic peptides have been linked recently.^7,8,24,54^ There have also been reports on the relation between fibril formation and environmental factors, such as oxidative stress and viral infections.^55–58^

Membranes constitute barriers and interfaces of complex composition and varying surface geometry.^40,59–62^ Along with sphingolipids, sterols, and membrane proteins, phospholipids are the major components of membranes that self-assemble into micelles, liposomes and bilayers (Figure 1c).^59^ Numerous studies identified a large impact of membrane composition on peptide self-assembly, ranging from acceleration to inhibition of the process.^17,39,71,72,63–70^ Here, we studied membrane compositions consisting of zwitterionic 1-palmitoyl-2-oleoyl-*sn*-glycero-3-phosphocholine (POPC) as a major lipid bilayer component (Figure 1d). Phosphatidylcholine (PC) lipids, such as POPC, are the main component of mammalian and bacterial cell membranes,^62^ and are typically used for biomimetic membrane studies.^73^ In addition to POPC, our model membranes and liposomes consisted of cholesterol, a typical and major component of mammalian membranes, as well as anionic 1-palmitoyl-2-oleoyl-*sn*-glycero-3-phospho-(1’-rac-glycerol) (POPG), a typical component of bacterial membranes.^8,74,75^ Furthermore, we studied the role of oxidative stress on peptide self-assembly by including the oxidized lipid 1-palmitoyl-2-azelaoyl-*sn*-glycero-3-phosphocholine (PazePC).^76–78^ Oxidative stress has previously been linked to ageing and neurodegenerative diseases.^79,80^ We used biophysical techniques to follow peptide self-assembly kinetics in the absence and presence of various lipid mixtures and peptide-to-lipid ratios to understand the impact on peptide secondary structure and peptide-membrane adsorption. Molecular dynamics (MD) simulations revealed molecular insights into the peptide-membrane interactions.

We observed differential effects of lipids on peptide self-assembly, depending on membrane composition and peptide-to-lipid ratio, with larger effects for the anionic POPG and the oxidized PazePC lipids, particularly for the U3.5 peptide. POPG and PazePC attracted the peptides onto their surface, driven by electrostatic interactions, and thereby influenced peptide secondary structure, leading to a large impact on peptide self-assembly.^81^ Interestingly, the same lipids and lipid mixtures, and peptide-to-lipid ratios led to differential effects for Aβ_40_ and U3.5 peptide, resulting from varying peptide-membrane attraction, hence secondary structure stabilization. Our results support the hypothesis that cellular changes, such as oxidative stress, could trigger peptide self-assembly processes and thus influence the onset and progression of related diseases.

## RESULTS AND DISCUSSION

To understand the influence of changes in the (cellular) membrane environment, namely the impact of an oxidized lipid, on the self-assembly of peptides into fibril structures, we initially performed experiments to follow the kinetics for the two peptides, amyloid β (1-40) (Aβ_40_) and uperin 3.5 (U3.5). We chose those peptides because of their similarity in forming fibrils as well as showing antimicrobial properties,^6,8,23,49^ but also their difference in overall charge, with Aβ_40_ being overall negatively charged and U3.5 positively charged (see Table 1). As membrane models, we used bilayers and liposomes or micelles (see Figure 1c) with the phospholipid POPC as the major component. Mixtures of POPC with cholesterol were used to model mammalian cells, mixtures of POPC with anionic POPG to mimic bacterial cells, and mixtures of POPC with PazePC to mimic oxidized membranes.^8,74,75^ PazePC has previously been identified as a major product of oxidative processes, and may be protonated or deprotonated at physiological pH (see Table 1).^76–78^

**Table 1.** Charges of the Peptides and Lipids at pH 7.4.

|  | **Charge**<br>**at pH**<br>**7.4** | **Charged Residues** |
| --- | --- | --- |
| **Peptides** |  |  |
| **Aβ<sub>40</sub>** | -2.9 | 6 x -1 (3 x Asp, 3 x Glu),<br>3 x +1 (Arg, 2 x Lys),<br>+0.1 (His), -1 (C-terminus),<br>+1 (N-terminus) |
| **U3.5** | +3 | -1 (Asp),<br>3 x +1 (Arg, 2 x Lys),<br>+1 (N-terminus) |
| **Lipids** |  |  |
| **POPC** | $\pm 0$ | -1 (phosphate),<br>+1 (choline) |
| **Cholesterol** | $\pm 0$ | neutral |
| **POPG** | -1 | -1 (phosphate) |
| **PazePC** | -1 to $\pm 0$ | -1 (phosphate), +1 (choline),<br>-1 (if azelaoyl carboxyl group<br>is deprotonated)<sup>78,82</sup> |

To probe the influence of the oxidized lipid PazePC on peptide self-assembly, we used thioflavin T (ThT) fluorescence assays. ThT is a commonly used dye to detect peptide self-assembly into amyloid fibrils, as it shows an enhanced fluorescence upon fibril binding.^83,84^ We studied the peptides Aβ_40_ and U3.5 without and with different amounts of lipids present (Figure 2). The peptide-to-lipid ratio was varied in order to study the situation with equal (molar) amounts of peptide and lipid (1:1) and excess of lipids (1:9). The lipids were added to the peptide solutions as liposomes, with the exception of POPC-PazePC and pure PazePC, which were present as micelles under the conditions used in this study, since PazePC has a relatively high critical micelle concentration (CMC) of ∼20 μM compared to POPC, POPG or cholesterol with CMC values in the nM range (see dynamic light scattering (DLS) measurements in SI Figure S2).^82,85–88^ The fluorescence assays consistently showed the greatest effects for ratios where lipids were in excess (shown in Figure 2, and SI Figure S3 for additional lipids and lipid mixtures). While Aβ_40_ aggregation was accelerated in the presence of all lipids (i.e. shorter lag times) (Figure 2a, c-e and SI Figure S3a, c, e, g), the aggregation of U3.5 was only minimally affected by POPC or cholesterol containing liposomes (POPC, cholesterol, POPC-cholesterol, 4:1) (Figure 2b, f and Si Figure S3b, h). If POPG or PazePC lipids were present (POPG, PazePC, POPC-POPG, 4:1, and POPC-PazePC, 7:3) (Figure 2b, g-h, SI Figure S3d, f), U3.5 aggregation was completely inhibited.

**Figure 2.**
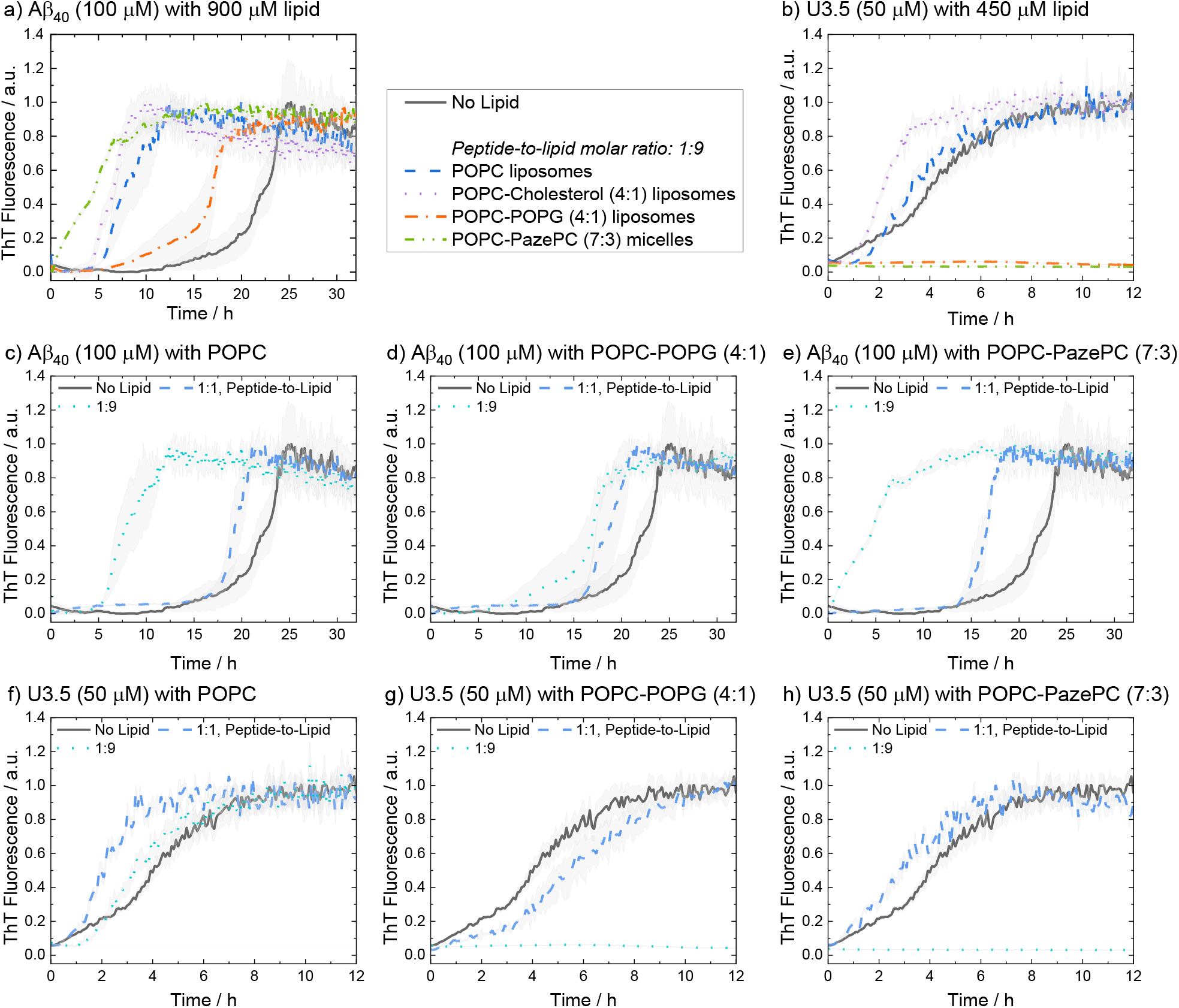
ThT fluorescence assays were performed to follow the kinetics of fibril formation. The peptides Aβ_40_ (100 μM) and U3.5 (50 μM) were studied in PBS buffer at pH 7.4 at 37°C. (a, b) Peptides were studied without and with different amounts of lipids present (peptide-to-lipid molar ratio: 1:1, 1:9). (c-h) The largest impact of the lipids on peptide aggregation was observed when lipid was added in excess (1:9). When peptide and lipid had the same concentration in the sample (1:1), smaller effects were observed. Data for pure POPC and lipid mixtures consisting of POPC-POPG (4:1) and POPC-PazePC (7:3) are shown. Additional data can be found in SI Figure S3. The data for U3.5 (b) without lipid present was previously reported and is included as a reference to all other lipids and Aβ_40_.^17^ The lines refer to the mean and the shadow areas to the SEM (standard error of the mean) of the replicates. Data were normalized to a maximum fluorescence of 1 (except in cases with inhibition of peptide aggregation).

The acceleration of Aβ_40_ aggregation in the presence of membrane surfaces is in agreement with previous studies^89^ while an enhanced β-sheet formation of the related Aβ_42_ peptide has been observed on oxidatively damaged surfaces, in particular.^69^ The inhibition of U3.5 aggregation in the presence of POPG containing liposomes (POPG, POPC-POPG, 4:1) is also in agreement with our prior work when DMPG (1,2-dimyristoyl-*sn*-glycero-3-phospho-(1’-rac-glycerol)) lipid systems were studied.^17^ In this work, we show that micelles containing the oxidized lipid PazePC have similar effects on peptide aggregation (Figure 2h and Si Figure S3f). Clearly, Aβ_40_ and U3.5 peptide aggregation were influenced by anionic lipids, albeit in a different direction, requiring us to consider the physicochemical properties of the peptides as illustrated above (Table 1).

If lipids were present at the same concentration as the peptide (1:1), Aβ_40_ aggregation was accelerated to a smaller extent compared to if lipids were present in an excess (1:9) (Figure 2c-e). For U3.5, lower lipid molar ratios (peptide-to-lipid ratio 1:1) had either no effect on peptide aggregation, or, interestingly, aggregation was accelerated, particularly when POPG liposomes were present (SI Figure S3d). When POPG or PazePC containing liposomes and micelles (POPG, PazePC, POPC-POPG, 4:1, and POPC-PazePC, 7:3) were present in excess (peptide-to-lipid molar ratio 1:9), U3.5 aggregation was completely inhibited (Figure 2b, g, h, SI Figure S3d,f). Liposomes and micelles present large surface-to-volume ratios in the nanometer size range (see SI Figure S2).^61,90,91^ In the presence of strong electrostatic surface attraction, the absorption of peptides onto micelles and liposomes can inhibit the formation of amyloid fibrils by depleting the free concentration of monomers in solution thereby reducing peptide mobility and flexibility. In the presence of weak surface attraction, micelle and liposome surfaces may act as potential adsorption and nucleation points for peptides and can seed aggregation.^8,39,92,93^ Similarly, if a small amount of surfaces with strong electrostatic attraction is present, surfaces can initiate the local concentration of peptide oligomers and thus accelerate peptide self-assembly.^17,90^ This may explain why U3.5 peptide aggregation was accelerated when POPG liposomes or PazePC micelles were present at low concentration (peptide-to-lipid molar ratio 1:1), but completely inhibited for ratios with lipid excess (1:9) (SI Figure S3d, f). The change in peptide-to-lipid ratio is also related a switch from peptide-rich to lipid-rich co-assemblies.^48,94^

Aβ_40_ peptide was influenced in a similar manner by all lipids and lipid mixtures and thus a similar adsorption mechanism of the peptide to the liposomes is expected. While Aβ_40_ has an overall negative charge, it has both positively and negatively charged side chains as well as many hydrophobic residues that may all provide potential points of attraction to membrane surfaces. In contrast, U3.5 peptide has an overall positive charge and thus attraction to negatively charged lipid headgroups is expected, leading to a strong influence of POPG and PazePC containing liposomes and micelles (POPG, PazePC, POPC-POPG, 4:1, and POPC-PazePC, 7.3). Uncharged POPC and cholesterol as well as POPC-cholesterol (4:1) liposomes did not significantly interact with U3.5 showing similar fluorescence profiles as without lipids present (Figure 2f, SI Figure S3b, h). While lower POPG liposome amounts (peptide-to-lipid molar ratio 1:1) may have provided a nucleation point leading to acceleration (Figure S3d), the inhibition of U3.5 aggregation at a peptide-to-lipid molar ratio of 1:9 was likely caused by trapping of all the U3.5 monomers at the micelle and liposome surfaces. Thus, these fluorescence results indicate a competition between oligomer seed formation and the inhibition of aggregation through the binding of available peptide monomers to the membrane surface.

To better understand the distinct effects of the membrane components on peptide self-assembly, we studied peptide secondary structure changes of Aβ_40_ and U3.5 using circular dichroism (CD) spectroscopy (Figure 3). The peptides were studied in the absence and presence of POPC, cholesterol, POPG and PazePC. The CD spectra show that the Aβ_40_ peptide aggregated and thus adapted a β-sheet conformation (λ_min_ at 215 nm) both in the absence and in the presence of lipids after two days of incubation. In contrast, U3.5 peptide showed β-sheet formation in the presence of POPC or cholesterol liposomes and without any lipid present (λ_min_ at 219 nm); however, the peptide was stabilized in an α-helical conformation (λ_min_ at 208 nm and 222 nm) when POPG liposomes or PazePC micelles were present;^95^ thus preventing β-sheet formation. The role of α-helical peptide conformations as potential intermediates toward fibrils and their high abundance at membrane surfaces has been discussed in the literature.^12,17,53^ It has previously been shown that peptides adopt a transmembrane conformation within membranes when the peptides are present in high concentration on the surface, resulting in an α-helical structure.^96^

**Figure 3.**
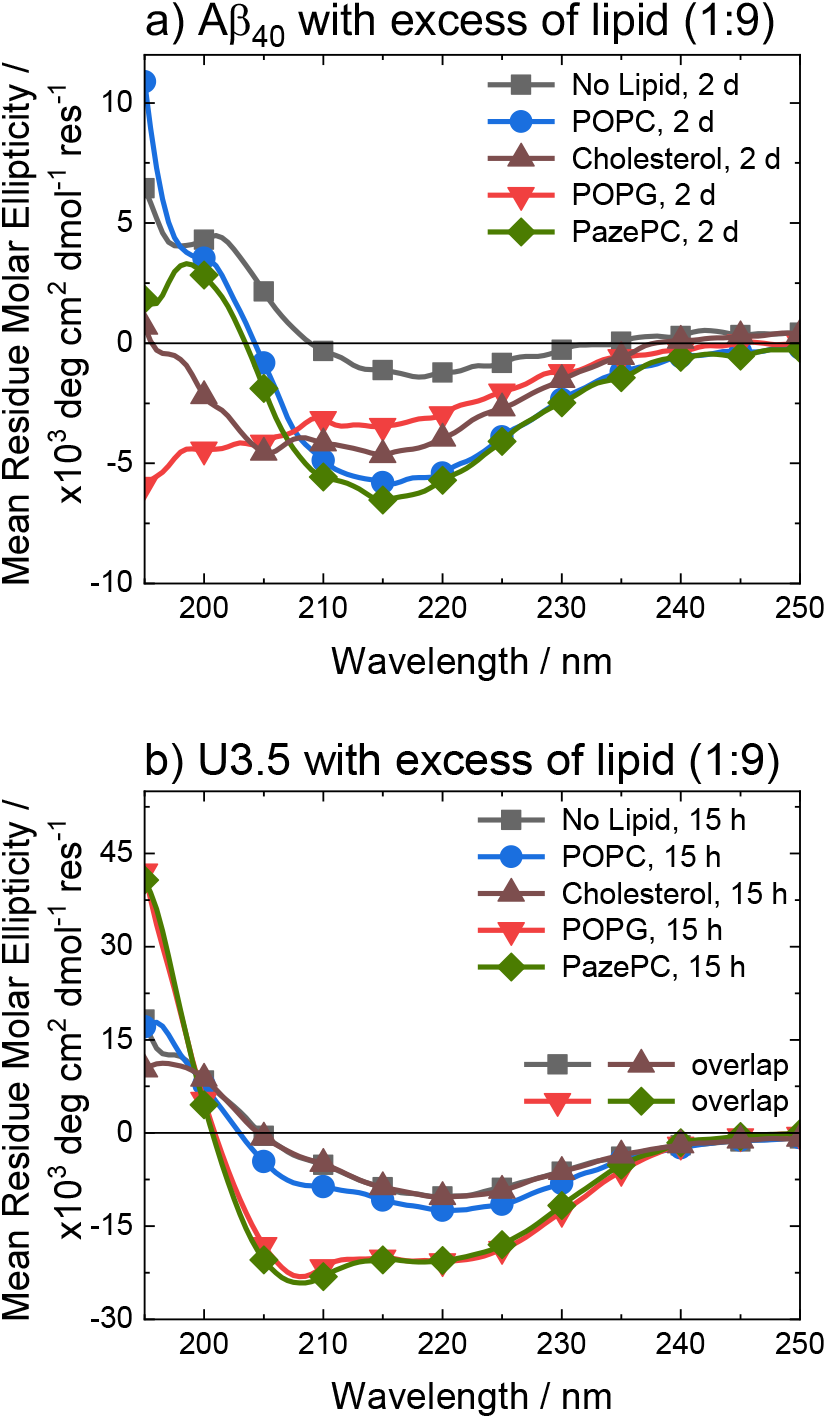
CD spectra of (a) Aβ_40_ and (b) U3.5 without and with excess of lipid (peptide-to-lipid molar ratio 1:9) in PBS buffer at pH 7.4 at 37°C. Aβ_40_ aggregation was studied at 100 μM with 900 μM of lipid present (for CD, it was diluted to 20 μM peptide and 180 μM lipid) and U3.5 peptide was studied at 50 μM with 450 μM lipid present. Samples were measured after 2 days or 15 hours, respectively. Note that the data for U3.5 (b) without lipid present and with cholesterol were previously reported and are included here as reference to all other lipids and Aβ_40_.^17^ Note that the symbols are used to distinguish the data sets and data were recorded every 0.5 nm.

Experiments with 2,2,2-trifluoroethanol (TFE) were performed to confirm that the solution environment has a distinct effect on Aβ_40_ secondary structure compared to U3.5. TFE is commonly used to enhance helical secondary structure.^17^ Our data show that while 40% TFE influenced the secondary structure of Aβ_40_ initially, the peptide aggregated after two days similarly to the Aβ_40_ samples with lipids (SI Figure S4). This is in contrast to U3.5 which was stabilized in its α-helical conformation for at least five days when 40% TFE was present, as previously reported^97^ (see Table S2 for quantitative secondary structure estimations from the experimental spectra). Bokvist *et al*. have shown that Aβ_40_ fibril formation is accelerated at membrane surfaces (for DMPC and DMPG) but prevented when the peptide was anchored in an α-helical conformation as a transmembrane peptide.^98^ In our study, we added liposomes and micelles to the peptides and thus an accelerating effect for Aβ_40_ was expected.

The stabilization of U3.5 in an α-helical conformation in the presence of excess POPG liposomes or PazePC micelles (Figure 3) is thus linked to the inhibitory effects on peptide aggregation observed in the ThT assays (SI Figure S3d, f), while an intermediate stabilization may accelerate peptide self-assembly. Since POPG and partially PazePC lipids are negatively charged and U3.5 positively charged, it seems that electrostatic attraction and thus strong adsorption of U3.5 to the membrane surface could be the cause for the inhibitory effects on fibril formation while stabilizing the peptide in an α-helical conformation. To probe differences in peptide-membrane binding, we performed quartz crystal microbalance (QCM) measurements (Figure 4). This technique enables monitoring of nanogram binding events to membrane surfaces using a surface-modified quartz crystal sensor.^99–102^

**Figure 4.**
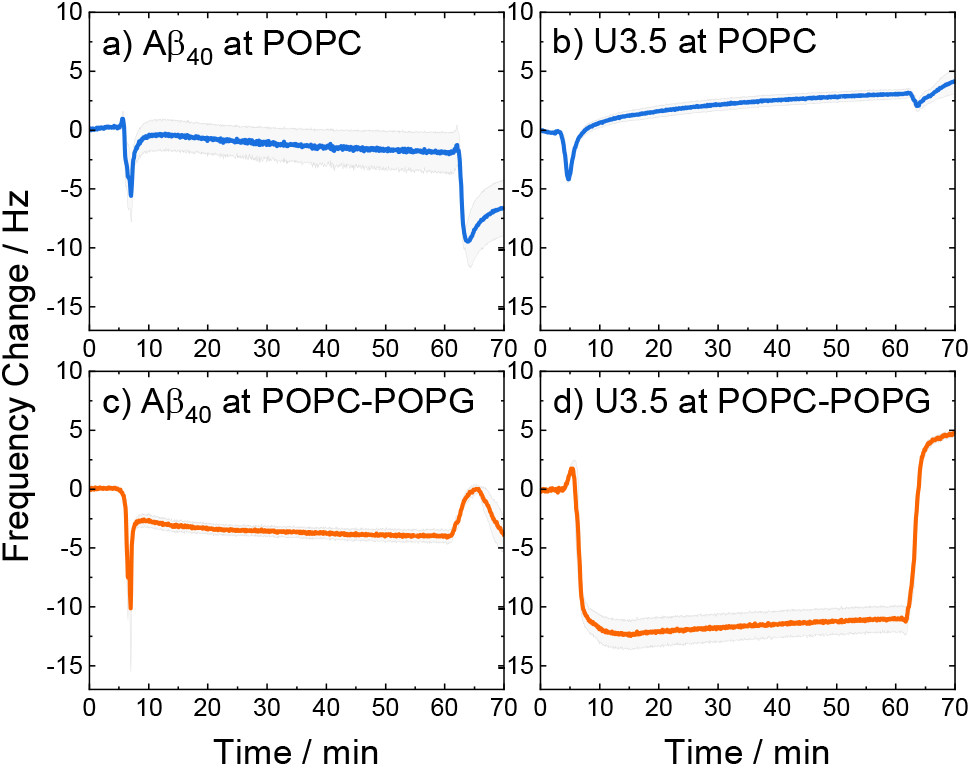
QCM changes in frequency of (a, c) Aβ_40_ and (b, d) U3.5 peptide (25 μM) interacting with (a, b) POPC and (c, d) POPC-POPG (4:1) lipid bilayers in PBS buffer at pH 7.4 at 22°C. The larger the negative change in frequency, the stronger the peptide mass binding to the membrane. A lipid bilayer is first deposited on the sensor surface before the peptide is introduced (0-15 min), kept incubating (15-60 min), and finally rinsed with buffer (60-70 min). The solid lines refer to the mean and the shadow areas to the SEM (standard error of the mean) of the replicates.

The QCM curves show a transient decrease in frequency and thus transient peptide binding for both peptides when interacting with POPC lipid bilayers (around 5 min), while only Aβ_40_ shows this behavior when interacting with POPC-POPG (4:1). In contrast, the U3.5 peptide remained bound within the POPC-POPG membrane over time until the measurement cell was rinsed with buffer. It can also be seen that both peptides showed greater mass binding with the POPC-POPG lipid bilayers (Figure 4c, d) compared to pure POPC (Figure 4a, b). While the peptide-membrane interaction mechanism requires a more detailed study for its elucidation, we used QCM measurements here for the purpose to probe differences in interaction affinity of the peptides to the membrane surfaces. Our results confirm the hypothesis that the peptides bind more strongly to negatively charged lipids, shown here for POPC-POPG due to its stability in liposome and membrane formation, and particularly true for the net positively charged U3.5 peptide.

Molecular dynamics (MD) simulations were performed to obtain detailed insight into peptide-membrane interactions.^78,103–105^ Both Aβ_40_ and U3.5 adsorbed to the lipid membranes within a few nanoseconds of simulation time. Representative snapshots of the most dominant structures of the simulations show a strong adsorption of particularly the U3.5 peptide to the membrane surface (Figure 5), while Aβ_40_ was less tightly bound. Aβ_40_ and U3.5 were studied with both an α-helical and unstructured (random) initial structure since many peptides are unstructured in solution and tend to form helices near membranes.^17,106^ Moreover, our atomistic MD simulations can only sample limited time scales and we thus considered both conformations as starting structures. Peptide-membrane adsorption appears particularly tight for the U3.5 peptide near POPC-POPG (4:1) and POPC-PazePC (7:3) membranes (see Figure 5 and SI Figures S5-S12 for simulation snapshots and mass density profiles). The tighter membrane binding of U3.5 compared to Aβ_40_ is consistent with our experimental QCM results (Figure 4) and is likely related to stronger electrostatic interactions, as it has been previously demonstrated for different charged surfaces.^97^ Analysis of the average distances between the phosphate headgroups of POPC in the outer membrane leaflet and the peptide C_α_ atoms confirmed the tighter binding of U3.5 peptide compared to Aβ_40_ to the membranes (Figure S13).

**Figure 5.**
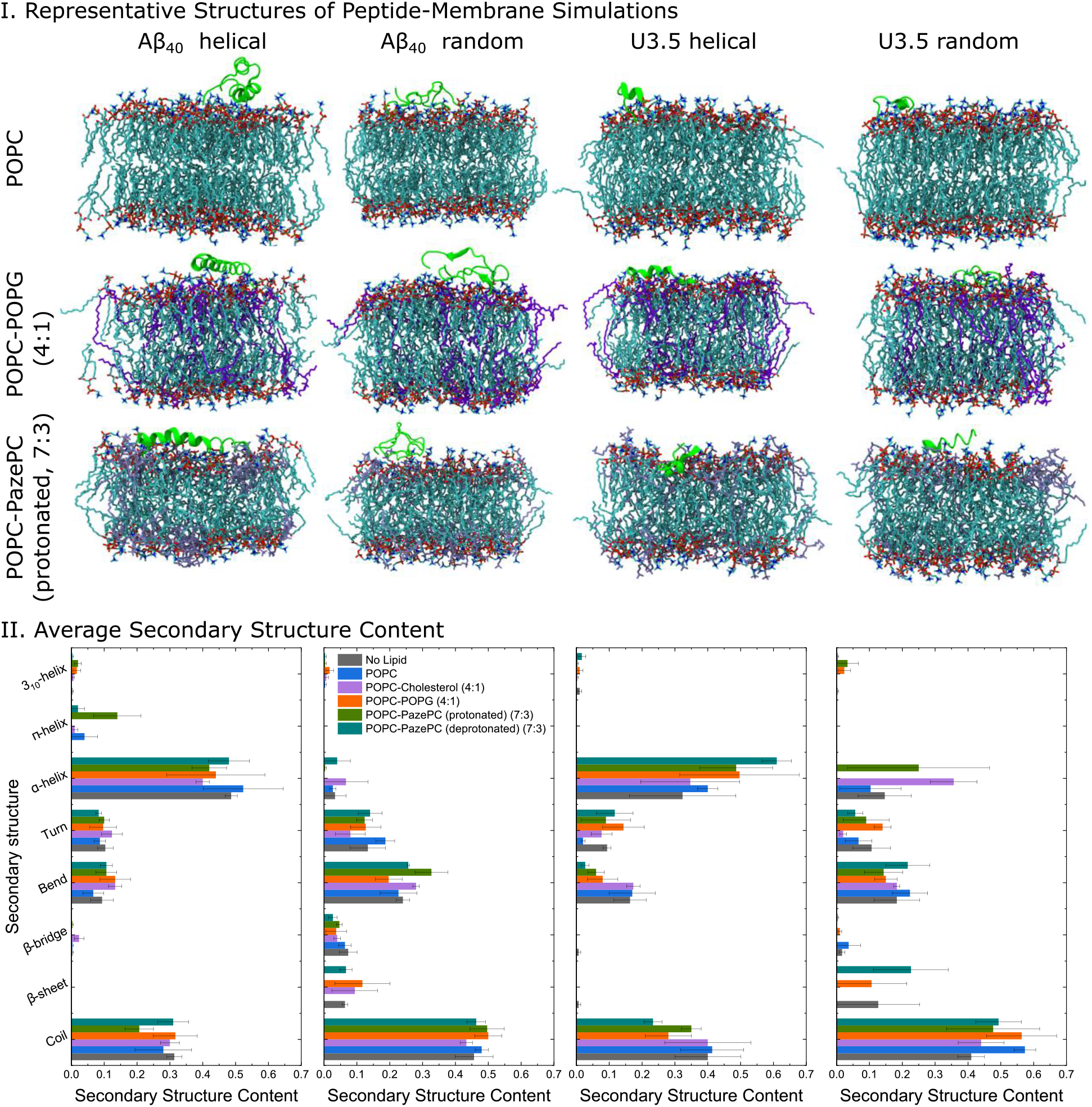
I) Representative structures of the peptide-membrane simulations at 303.15 K and with 0.15 mM NaCl in water. The central structure of the largest structural cluster during the last 10 ns simulation time of all replicates for each peptide is shown when studied near POPC, POPC-POPG (4:1), and POPC-PazePC (protonated) (7:3) membranes. II) The average secondary structure content of the peptides during the last 10 ns simulation time of all replicates is shown. MD simulation snapshots were visualized in VMD 1.93.^109^

When analyzing the secondary structure of the peptide conformations in the simulations, we observed a greater β-sheet/bridge formation when the initial peptide structure was random coil, while the α-helical starting structure remained more stable overall. Significantly, our MD simulations confirmed a strong α-helix stabilization for the U3.5 peptide (helical starting structure) in the presence of POPC-POPG (4:1) and POPC-PazePC (7:3) membranes (see Figure 5, secondary structure content for U3.5 helical). Note that we studied membranes containing both protonated as well as deprotonated PazePC due to the potential presence of both under experimental conditions (pH 7.4). This agrees with our observation that fibril formation was completely inhibited for U3.5 peptide when POPG or PazePC containing membranes were present in excess (Figure 2). In contrast, Aβ_40_ secondary structure was not significantly influenced by the presence of the membranes, confirming our CD spectroscopy experiments (Figure 3), where we observed aggregation into β-sheet rich structures for Aβ_40_ in all cases. The MD simulations are in overall agreement with our experimental observations with a tighter membrane binding and secondary structure stabilization (see Figure 5) and thus larger influence on peptide aggregation for the U3.5 peptide. The relevance of the environmental conditions on the U3.5 peptide conformation has recently also been reported by Landau *et al*. who have determined an cross-α helical structure when in an environment of a polyether based on polypropylene glycol,^12^ and a cross-β fibril structure in aqueous solution.^107^ Previous studies also found that the related Aβ_42_ peptide is influenced in oligomer structure formation depending on the membrane composition and its environment.^108^

Analysis of the closest interactions between the peptides and the membrane components confirmed our suggestion that differences in the charge between both peptides caused the differential impact caused by POPC, anionic POPG and oxidized PazePC lipids (Figure 6). The cationic amino acids arginine and lysine in positions 5, 16 and 28 in Aβ_40_, and in positions 7, 8 and 14 in U3.5 showed minima in peptide-membrane distance, and thus indicate the most dominant peptide-membrane interactions. Electrostatic attraction was particularly observed for the U3.5 peptide. In previous work, we already demonstrated the high relevance of position 7 (arginine) in U3.5 for peptide aggregation and membrane interactions.^8,17^ Our MD simulations show that positions 8 and 14 were of high relevance for the initial membrane interactions for the U3.5 peptide, for both the simulations with an α-helical and random starting structure. We note that longer simulation time scales would be required to study more detailed effects on the membrane integrity. However, our study here focused on the effects on peptide adsorption and self-assembly.

**Figure 6.**
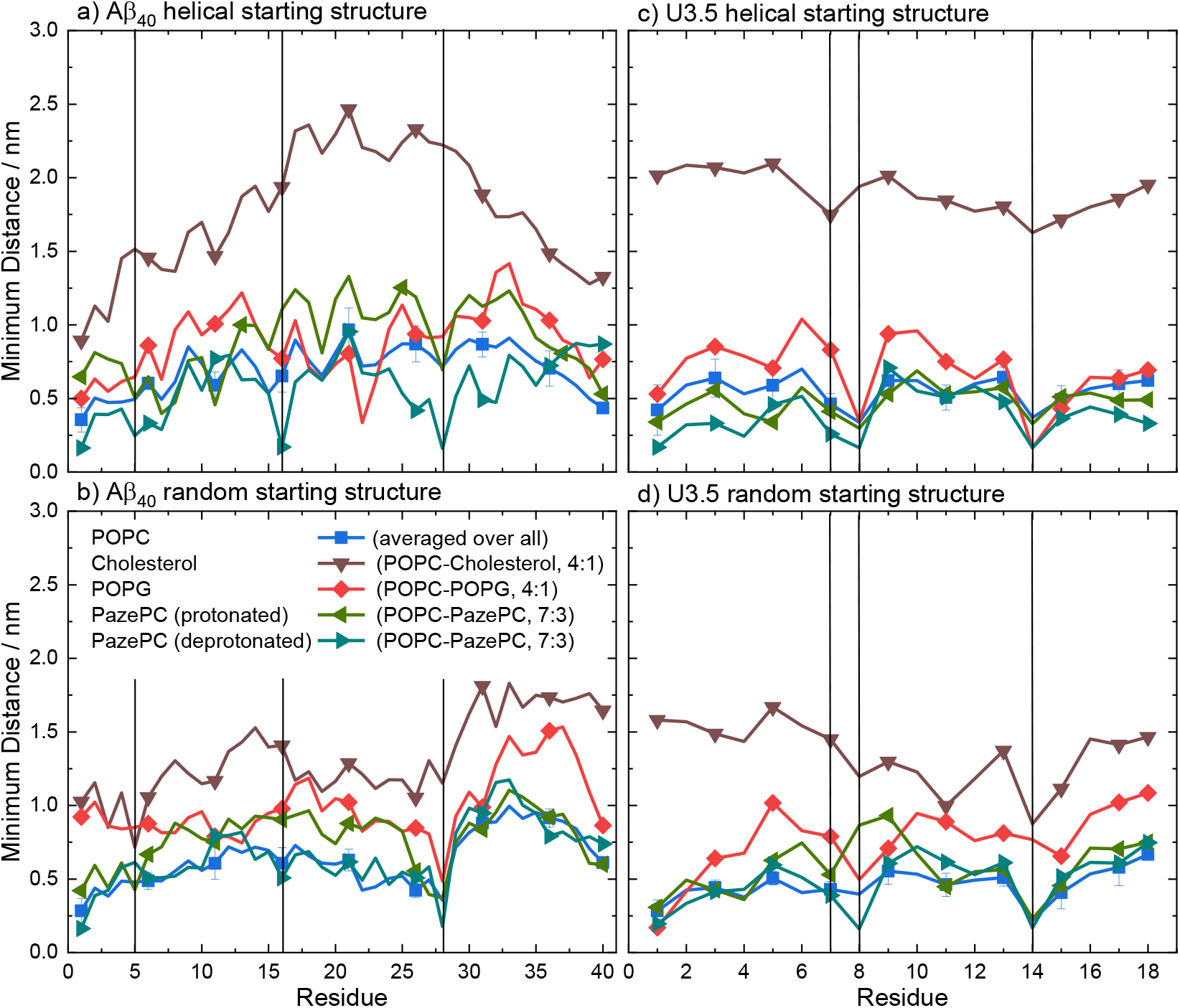
Average minimum distances between the peptides (a, b) Aβ_40_ and (c, d) U3.5 and the lipid bilayer components POPC, cholesterol, POPG, and PazePC during the last 10 ns simulation time of all replicates. The vertical lines at residues 5 (arginine), 16 (lysine) and 28 (lysine) for Aβ_40_ and at residues 7 (arginine), 8 (arginine) and 14 (lysine) for U3.5 indicate the positively charged residues in both peptides to guide identifying the closest peptide-membrane interactions. Note that the symbols are used to distinguish the data sets and each residue has a data point.

In summary, our data have shown that membrane lipid composition modulates the effect of membranes on peptide self-assembly. For the overall negatively charged and highly hydrophobic Aβ_40_ peptide, peptide aggregation was accelerated by all membranes at the peptide-to-lipid ratios that we tested, especially by POPC and PazePC containing membranes. In contrast, the net positively charged U3.5 peptide was not significantly influenced by POPC or cholesterol but showed some acceleration at low amounts of POPG and PazePC containing membranes, and complete inhibition when POPG and PazePC lipids were present in excess. Our QCM experiments and MD simulations confirmed a strong binding of U3.5 to POPG and PazePC containing membranes, with stabilization of the peptide in the α-helical conformation, revealed by CD spectroscopy. The impact of membrane lipids on peptide aggregation, as well as changes in membrane composition, must therefore be considered to be peptide-specific. The peptide sequence as well as the peptide aggregation propensity are important factors next to the peptide-membrane surface attraction.^110^ The membrane composition, influenced by oxidation processes, results in distinct physical properties of the membranes,^77,111–114^ which in turn impact the effects on peptide aggregation.^115^ In our study, we have investigated the effects for the net negatively charged Aβ_40_ peptide and the net positively charged U3.5 peptide. Electrostatic interactions mediated by the presence of charged peptide residues were highly relevant for the U3.5 peptide, while Aβ_40_ peptide experienced intermediate affinity to all membranes. This is in agreement with previous work that identified electrostatic interactions to be particularly important for cationic peptides and hydrophobic peptide residues to drive the interactions of nonpolar residues with membranes.^66,116^ The sequence and number of positively charged residues in peptides, such as arginine and lysine, may thus have important influence on the degree and direction in which anionic and oxidized lipids affect peptide aggregation.

## CONCLUSIONS

This work suggests the importance of (cellular) lipid membranes, particularly their biochemical composition, for biologically relevant processes, such as peptide adsorption, secondary structure stabilization and aggregation. Changes in the membrane structure due to oxidative reactions or the infection with bacteria and thus the exposure to new and distinctly modified cell surfaces may lead to enhanced peptide adsorption to membrane surfaces, resulting in the acceleration or inhibition of peptide aggregation, and in turn amyloid fibril formation as shown in our work. It seems obvious that these effects may have implications for the link between amyloidogenic peptides and the development of neurodegenerative diseases. The strong effect of the anionic, bacteria-mimicking, and oxidized membrane lipids on the antimicrobial peptide U3.5 emphasize our hypotheses about existing links between antimicrobial and amyloidogenic peptides, as well as to infection and (oxidative) stress. Both Aβ_40_ and U3.5 may have a functional role in organisms and their self-assembly into α-helical or β-sheet rich conformations be linked to functional or disease-related states. Changes in the cellular membrane and peptide conformation due to stress could thus trigger functional loss of peptides and proteins.

## Author Contributions

The study was designed and conceptualized by TJ, BA, HJR and LLM. Biophysical studies were performed and analyzed by TJ (ThT fluorescence assays, CD spectroscopy, DLS measurements), SP (QCM and CD measurements) and TD (CD spectroscopy). Computer simulations were performed and analyzed by TJ. The results were discussed and interpreted by TJ, BA, HRJ and LLM. The manuscript was written by TJ and advanced by all authors.

## Notes

The authors declare no competing financial interest.

## ACKNOWLEDGMENT

The authors thank the members of the DFG SFB-TRR 102, particularly Prof. Daniel Huster, for interesting and stimulating discussions. This work was funded by the Deutsche Forschungsgemeinschaft (DFG, German Research Foundation, project number 189853844, SFB-TRR 102, B1). TJ thanks the Friedrich-Ebert-Stiftung for a PhD fellowship, and the Australian Government, Department of Education and Training, and Scope Global for the support through an Endeavour Research Fellowship.

## ABBREVIATIONS

Aβ_40_: amyloid β (1-40)
AMP: antimicrobial peptide
CD: circular dichroism
CMC: critical micelle concentration
DLS: dynamic light scattering
DMPG: 1,2-dimyristoyl-*sn*-glycero-3-phospho-(1’-rac-glycerol)
MD: molecular dynamics
PazePC: 1-palmitoyl-2-azelaoyl-*sn*-glycero-3-phosphocholine
PBS: phosphate-buffered saline: POPC 1-palmitoyl-2-oleoyl-*sn*-glycero-3-phosphocholine
POPG: 1-palmitoyl-2-oleoyl-*sn*-glycero-3-phospho-(1’-rac-glycerol)
QCM: quartz crystal microbalance
TFE: 2,2,2-trifluoroethanol
ThT: thioflavin T; U3.5, uperin 3.5

## Notes

### Competing Interest Statement

The authors have declared no competing interest.

## REFERENCES

(1) Chiti, F.; Dobson, C. M. Protein Misfolding, Amyloid Formation, and Human Disease: A Summary of Progress Over the Last Decade. Annu. Rev. Biochem. 2017, 86 (1), 27–68. https://doi.org/10.1146/annurev-biochem-061516-045115.

(2) Glenner, G. G.; Wong, C. W. Alzheimer’s Disease: Initial Report of the Purification and Characterization of a Novel Cerebrovascular Amyloid Protein. Biochem. Biophys. Res. Commun. 1984, 120 (3), 885–890. https://doi.org/10.1016/S0006-291X(84)80190-4.

(3) Goedert, M.; Spillantini, M. G. A Century of Alzheimer’s Disease. Science 2006, 314 (5800), 777–781. https://doi.org/10.1126/science.1132814.

(4) Chiti, F.; Dobson, C. M. Protein Misfolding, Functional Amyloid, and Human Disease. Annu. Rev. Biochem. 2006, 75 (1), 333–366. https://doi.org/10.1146/annurev.biochem.75.101304.123901.

(5) Wei, G.; Su, Z.; Reynolds, N. P.; Arosio, P.; Hamley, I. W.; Gazit, E.; Mezzenga, R. Self-Assembling Peptide and Protein Amyloids: From Structure to Tailored Function in Nanotechnology. Chem. Soc. Rev. 2017, 46 (15), 4661–4708. https://doi.org/10.1039/C6CS00542J.

(6) Bradford, A.; Bowie, J.; Tyler, M.; Wallace, J. New Antibiotic Uperin Peptides From the Dorsal Glands of the Australian Toadlet Uperoleia Mjobergii. Aust. J. Chem. 1996, 49 (12), 1325–1331. https://doi.org/10.1071/CH9961325.

(7) Calabrese, A. N.; Liu, Y.; Wang, T.; Musgrave, I. F.; Pukala, T. L.; Tabor, R. F.; Martin, L. L.; Carver, J. A.; Bowie, J. H. The Amyloid Fibril-Forming Properties of the Amphibian Antimicrobial Peptide Uperin 3.5. ChemBioChem 2016, 17 (3), 239–246. https://doi.org/10.1002/cbic.201500518.

(8) Martin, L. L.; Kubeil, C.; Piantavigna, S.; Tikkoo, T.; Gray, N. P.; John, T.; Calabrese, A. N.; Liu, Y.; Hong, Y.; Hossain, M. A.; Patil, N.; Abel, B.; Hoffmann, R.; Bowie, J. H.; Carver, J. A. Amyloid Aggregation and Membrane Activity of the Antimicrobial Peptide Uperin 3.5. Pept. Sci. 2018, 110 (3), e24052. https://doi.org/10.1002/pep2.24052.

(9) Nelson, R.; Sawaya, M. R.; Balbirnie, M.; Madsen, A. Ø.; Riekel, C.; Grothe, R.; Eisenberg, D. Structure of the Cross-β Spine of Amyloid-like Fibrils. Nature 2005, 435 (7043), 773– 778. https://doi.org/10.1038/nature03680.

(10) Makin, O. S.; Serpell, L. C. Structures for Amyloid Fibrils. FEBS J. 2005, 272 (23), 5950–5961. https://doi.org/10.1111/j.1742-4658.2005.05025.x.

(11) Tayeb-Fligelman, E.; Tabachnikov, O.; Moshe, A.; Goldshmidt-Tran, O.; Sawaya, M. R.; Coquelle, N.; Colletier, J.; Landau, M. The Cytotoxic Staphylococcus Aureus PSMα3 Reveals a Cross-α Amyloid-like Fibril. Science 2017, 355 (6327), 831–833. https://doi.org/10.1126/science.aaf4901.

(12) Salinas, N.; Tayeb-Fligelman, E.; Sammito, M. D.; Bloch, D.; Jelinek, R.; Noy, D.; Usón, I.; Landau, M. The Amphibian Antimicrobial Peptide Uperin 3.5 Is a Cross-α/Cross-β Chameleon Functional Amyloid. Proc. Natl. Acad. Sci. 2021, 118 (3), e2014442118. https://doi.org/10.1073/pnas.2014442118.

(13) Engelberg, Y.; Landau, M. The Human LL-37(17-29) Antimicrobial Peptide Reveals a Functional Supramolecular Structure. Nat. Commun. 2020, 11 (1), 1–10. https://doi.org/10.1038/s41467-020-17736-x.

(14) Ragonis-Bachar, P.; Rayan, B.; Barnea, E.; Engelberg, Y.; Upcher, A.; Landau, M. Natural Antimicrobial Peptides Self-Assemble as α/β Chameleon Amyloids. bioRxiv 2022, 1–67. https://doi.org/10.1101/2022.06.23.497336.

(15) Cohen, S. I. A.; Vendruscolo, M.; Dobson, C. M.; Knowles, T. P. J. From Macroscopic Measurements to Microscopic Mechanisms of Protein Aggregation. J. Mol. Biol. 2012, 421 (2–3), 160–171. https://doi.org/10.1016/j.jmb.2012.02.031.

(16) Meisl, G.; Kirkegaard, J. B.; Arosio, P.; Michaels, T. C. T.; Vendruscolo, M.; Dobson, C. M.; Linse, S.; Knowles, T. P. J. Molecular Mechanisms of Protein Aggregation from Global Fitting of Kinetic Models. Nat. Protoc. 2016, 11 (2), 252–272. https://doi.org/10.1038/nprot.2016.010.

(17) John, T.; Dealey, T. J. A.; Gray, N. P.; Patil, N. A.; Hossain, M. A.; Abel, B.; Carver, J. A.; Hong, Y.; Martin, L. L. The Kinetics of Amyloid Fibrillar Aggregation of Uperin 3.5 Is Directed by the Peptide’s Secondary Structure. Biochemistry 2019, 58 (35), 3656–3668. https://doi.org/10.1021/acs.biochem.9b00536.

(18) Fezoui, Y.; Teplow, D. B. Kinetic Studies of Amyloid β-Protein Fibril Assembly: DIFFERENTIAL EFFECTS OF α-HELIX STABILIZATION. J. Biol. Chem. 2002, 277 (40), 36948–36954. https://doi.org/10.1074/jbc.M204168200.

(19) De Carufel, C. A.; Quittot, N.; Nguyen, P. T.; Bourgault, S. Delineating the Role of Helical Intermediates in Natively Unfolded Polypeptide Amyloid Assembly and Cytotoxicity. Angew. Chemie - Int. Ed. 2015, 54 (48), 14383–14387. https://doi.org/10.1002/anie.201507092.

(20) Ray, S.; Holden, S.; Prasad, A. K.; Martin, L. L.; Panwar, A. S. Exploring the Role of Peptide Helical Stability in the Propensity of Uperin 3.x Peptides toward Beta-Aggregation. J. Phys. Chem. B 2020, 124 (51), 11659–11670. https://doi.org/10.1021/acs.jpcb.0c10000.

(21) Brothers, H. M.; Gosztyla, M. L.; Robinson, S. R. The Physiological Roles of Amyloid-β Peptide Hint at New Ways to Treat Alzheimer’s Disease. Front. Aging Neurosci. 2018, 10 (APR), 1–16. https://doi.org/10.3389/fnagi.2018.00118.

(22) Fowler, D. M.; Koulov, A. V.; Balch, W. E.; Kelly, J. W. Functional Amyloid - from Bacteria to Humans. Trends Biochem. Sci. 2007, 32 (5), 217–224. https://doi.org/10.1016/j.tibs.2007.03.003.

(23) Soscia, S. J.; Kirby, J. E.; Washicosky, K. J.; Tucker, S. M.; Ingelsson, M.; Hyman, B.; Burton, M. A.; Goldstein, L. E.; Duong, S.; Tanzi, R. E.; Moir, R. D. The Alzheimer’s Disease-Associated Amyloid β-Protein Is an Antimicrobial Peptide. PLoS One 2010, 5 (3). https://doi.org/10.1371/journal.pone.0009505.

(24) Caillon, L.; Killian, J. A.; Lequin, O.; Khemtémourian, L. Biophysical Investigation of the Membrane-Disrupting Mechanism of the Antimicrobial and Amyloid-Like Peptide Dermaseptin S9. PLoS One 2013, 8 (10), e75528. https://doi.org/10.1371/journal.pone.0075528.

(25) Harris, F.; Dennison, S. R.; Phoenix, D. A. Aberrant Action of Amyloidogenic Host Defense Peptides: A New Paradigm to Investigate Neurodegenerative Disorders? FASEB J. 2012, 26 (5), 1776–1781. https://doi.org/10.1096/fj.11-199208.

(26) Ruggeri, F.; Zhang, F.; Lind, T.; Bruce, E. D.; Lau, B. L. T.; Cárdenas, M. Non-Specific Interactions between Soluble Proteins and Lipids Induce Irreversible Changes in the Properties of Lipid Bilayers. Soft Matter 2013, 9 (16), 4219–4226. https://doi.org/10.1039/C3SM27769K.

(27) Magno, A.; Caflisch, A.; Pellarin, R. Crowding Effects on Amyloid Aggregation Kinetics. J. Phys. Chem. Lett. 2010, 1 (20), 3027–3032. https://doi.org/10.1021/jz100967z.

(28) White, D. A.; Buell, A. K.; Knowles, T. P. J.; Welland, M. E.; Dobson, C. M. Protein Aggregation in Crowded Environments. J. Am. Chem. Soc. 2010, 132 (14), 5170–5175. https://doi.org/10.1021/ja909997e.

(29) Yeung, P. S. W.; Axelsen, P. H. The Crowded Environment of a Reverse Micelle Induces the Formation of β-Strand Seed Structures for Nucleating Amyloid Fibril Formation. J. Am. Chem. Soc. 2012, 134 (14), 6061–6063. https://doi.org/10.1021/ja3004478.

(30) Stefani, M. Biochemical and Biophysical Features of Both Oligomer/Fibril and Cell Membrane in Amyloid Cytotoxicity. FEBS J. 2010, 277 (22), 4602–4613. https://doi.org/10.1111/j.1742-4658.2010.07889.x.

(31) Cao, P.; Abedini, A.; Wang, H.; Tu, L. H.; Zhang, X.; Schmidt, A. M.; Raleigh, D. P. Islet Amyloid Polypeptide Toxicity and Membrane Interactions. Proc. Natl. Acad. Sci. U. S. A. 2013, 110 (48), 19279–19284. https://doi.org/10.1073/pnas.1305517110.

(32) Williams, T. L.; Serpell, L. C. Membrane and Surface Interactions of Alzheimer’s Aβ Peptide - Insights into the Mechanism of Cytotoxicity. FEBS J. 2011, 278 (20), 3905–3917. https://doi.org/10.1111/j.1742-4658.2011.08228.x.

(33) Axelsen, P. H.; Komatsu, H.; Murray, I. V. J. Oxidative Stress and Cell Membranes in the Pathogenesis of Alzheimer’s Disease. Physiology (Bethesda). 2011, 26 (1), 54–69. https://doi.org/10.1152/physiol.00024.2010.

(34) Kotler, S. A.; Walsh, P.; Brender, J. R.; Ramamoorthy, A. Differences between Amyloid-β Aggregation in Solution and on the Membrane: Insights into Elucidation of the Mechanistic Details of Alzheimer’s Disease. Chem. Soc. Rev. 2014, 43 (19), 6692–6700. https://doi.org/10.1039/C3CS60431D.

(35) Butterfield, S. M.; Lashuel, H. A. Amyloidogenic Protein-Membrane Interactions: Mechanistic Insight from Model Systems. Angew. Chemie Int. Ed. 2010, 49 (33), 5628–5654. https://doi.org/10.1002/anie.200906670.

(36) Rawat, A.; Langen, R.; Varkey, J. Membranes as Modulators of Amyloid Protein Misfolding and Target of Toxicity. Biochim. Biophys. Acta - Biomembr. 2018, 1860 (9), 1863–1875. https://doi.org/10.1016/j.bbamem.2018.04.011.

(37) Bandyopadhyay, A.; Sannigrahi, A.; Chattopadhyay, K. Membrane Composition and Lipid to Protein Ratio Modulate Amyloid Kinetics of Yeast Prion Protein. RSC Chem. Biol. 2021, 2 (2), 592–605. https://doi.org/10.1039/D0CB00203H.

(38) Han, X.; Wu, X.; Lv, L.; Li, C. Inhibiting and Catalysing Amyloid Fibrillation at Dynamic Lipid Interfaces. J. Colloid Interface Sci. 2019, 543, 256–262. https://doi.org/10.1016/j.jcis.2019.02.072.

(39) Krausser, J.; Knowles, T. P. J.; Šarić, A. Physical Mechanisms of Amyloid Nucleation on Fluid Membranes. Proc. Natl. Acad. Sci. U. S. A. 2020, 117 (52), 33090–33098. https://doi.org/10.1073/pnas.2007694117.

(40) Sanguanini, M.; Baumann, K. N.; Preet, S.; Chia, S.; Habchi, J.; Knowles, T. P. J.; Vendruscolo, M. Complexity in Lipid Membrane Composition Induces Resilience to Aβ 42 Aggregation. ACS Chem. Neurosci. 2020, 11 (9), 1347–1352. https://doi.org/10.1021/acschemneuro.0c00101.

(41) Gorbenko, G. P.; Kinnunen, P. K. J. The Role of Lipid-Protein Interactions in Amyloid-Type Protein Fibril Formation. Chem. Phys. Lipids 2006, 141 (1–2), 72–82. https://doi.org/10.1016/j.chemphyslip.2006.02.006.

(42) Press-Sandler, O.; Miller, Y. Molecular Insights into the Primary Nucleation of Polymorphic Amyloid β Dimers in DOPC Lipid Bilayer Membrane. Protein Sci. 2022, 31 (5), 1–14. https://doi.org/10.1002/pro.4283.

(43) Burke, K. A.; Yates, E. A.; Legleiter, J. Biophysical Insights into How Surfaces, Including Lipid Membranes, Modulate Protein Aggregation Related to Neurodegeneration. Front. Neurol. 2013, 4 MAR (March), 1–17. https://doi.org/10.3389/fneur.2013.00017.

(44) Press-Sandler, O.; Miller, Y. Molecular Mechanisms of Membrane-Associated Amyloid Aggregation: Computational Perspective and Challenges. Biochim. Biophys. Acta - Biomembr. 2018, 1860 (9), 1889–1905. https://doi.org/10.1016/j.bbamem.2018.03.014.

(45) Bode, D. C.; Freeley, M.; Nield, J.; Palma, M.; Viles, J. H. Amyloid-β Oligomers Have a Profound Detergent-like Effect on Lipid Membrane Bilayers, Imaged by Atomic Force and Electron Microscopy. J. Biol. Chem. 2019, 294 (19), 7566–7572. https://doi.org/10.1074/jbc.AC118.007195.

(46) Martel, A.; Antony, L.; Gerelli, Y.; Porcar, L.; Fluitt, A.; Hoffmann, K.; Kiesel, I.; Vivaudou, M.; Fragneto, G.; De Pablo, J. J. Membrane Permeation versus Amyloidogenicity: A Multitechnique Study of Islet Amyloid Polypeptide Interaction with Model Membranes. J. Am. Chem. Soc. 2017, 139 (1), 137–148. https://doi.org/10.1021/jacs.6b06985.

(47) Andrade, S.; Loureiro, J. A.; Pereira, M. C. The Role of Amyloid B-Biomembrane Interactions in the Pathogenesis of Alzheimer’s Disease: Insights from Liposomes as Membrane Models. ChemPhysChem 2021, 1–20. https://doi.org/10.1002/cphc.202100124.

(48) Sparr, E.; Linse, S. Lipid-Protein Interactions in Amyloid Formation. Biochim. Biophys. Acta - Proteins Proteomics 2019, 1867 (5), 455–457. https://doi.org/10.1016/j.bbapap.2019.03.006.

(49) Li, Y.; Tang, H.; Andrikopoulos, N.; Javed, I.; Cecchetto, L.; Nandakumar, A.; Kakinen, A.; Davis, T. P.; Ding, F.; Ke, P. C. The Membrane Axis of Alzheimer’s Nanomedicine. Adv. NanoBiomed Res. 2021, 1 (1), 2000040. https://doi.org/10.1002/anbr.202000040.

(50) Shai, Y. Mechanism of the Binding, Insertion and Destabilization of Phospholipid Bilayer Membranes by α-Helical Antimicrobial and Cell Non-Selective Membrane-Lytic Peptides. Biochim. Biophys. Acta - Biomembr. 1999, 1462 (1–2), 55–70. https://doi.org/10.1016/S0005-2736(99)00200-X.

(51) Nguyen, L. T.; Haney, E. F.; Vogel, H. J. The Expanding Scope of Antimicrobial Peptide Structures and Their Modes of Action. Trends Biotechnol. 2011, 29 (9), 464–472. https://doi.org/10.1016/j.tibtech.2011.05.001.

(52) Zheng, Q.; Carty, S. N.; Lazo, N. D. Helix Dipole and Membrane Electrostatics Delineate Conformational Transitions in the Self-Assembly of Amyloidogenic Peptides. Langmuir 2020, 36 (29), 8389–8397. https://doi.org/10.1021/acs.langmuir.0c00723.

(53) Prasad, A. K.; Tiwari, C.; Ray, S.; Holden, S.; Armstrong, D. A.; Rosengren, K. J.; Rodger, A.; Panwar, A. S.; Martin, L. L. Secondary Structure Transitions for a Family of Amyloidogenic, Antimicrobial Uperin 3 Peptides in Contact with Sodium Dodecyl Sulfate. Chempluschem 2022, 87 (1), e202100408. https://doi.org/10.1002/cplu.202100408.

(54) Last, N. B.; Miranker, A. D. Common Mechanism Unites Membrane Poration by Amyloid and Antimicrobial Peptides. Proc. Natl. Acad. Sci. U. S. A. 2013, 110 (16), 6382–6387. https://doi.org/10.1073/pnas.1219059110.

(55) Cheignon, C.; Tomas, M.; Bonnefont-Rousselot, D.; Faller, P.; Hureau, C.; Collin, F. Oxidative Stress and the Amyloid Beta Peptide in Alzheimer’s Disease. Redox Biol. 2018, 14 (September 2017), 450–464. https://doi.org/10.1016/j.redox.2017.10.014.

(56) Eimer, W. A.; Vijaya Kumar, D. K.; Navalpur Shanmugam, N. K.; Rodriguez, A. S.; Mitchell, T.; Washicosky, K. J.; György, B.; Breakefield, X. O.; Tanzi, R. E.; Moir, R. D. Alzheimer’s Disease-Associated β-Amyloid Is Rapidly Seeded by Herpesviridae to Protect against Brain Infection. Neuron 2018, 99 (1), 56-63.e3. https://doi.org/10.1016/j.neuron.2018.06.030.

(57) Cutler, R. G.; Kelly, J.; Storie, K.; Pedersen, W. A.; Tammara, A.; Hatanpaa, K.; Troncoso, J. C.; Mattson, M. P. Involvement of Oxidative Stress-Induced Abnormalities in Ceramide and Cholesterol Metabolism in Brain Aging and Alzheimer’s Disease. Proc. Natl. Acad. Sci. U. S. A. 2004, 101 (7), 2070–2075. https://doi.org/10.1073/pnas.0305799101.

(58) Ezzat, K.; Pernemalm, M.; Pålsson, S.; Roberts, T. C.; Järver, P.; Dondalska, A.; Bestas, B.; Sobkowiak, M. J.; Levänen, B.; Sköld, M.; et al. The Viral Protein Corona Directs Viral Pathogenesis and Amyloid Aggregation. Nat. Commun. 2019, 10 (1), 2331. https://doi.org/10.1038/s41467-019-10192-2.

(59) O’Brien, J. S. Cell Membranes—Composition: Structure: Function. J. Theor. Biol. 1967, 15 (3), 307–324. https://doi.org/10.1016/0022-5193(67)90140-3.

(60) Kinoshita, M.; Kakimoto, E.; Terakawa, M. S.; Lin, Y.; Ikenoue, T.; So, M.; Sugiki, T.; Ramamoorthy, A.; Goto, Y.; Lee, Y. H. Model Membrane Size-Dependent Amyloidogenesis of Alzheimer’s Amyloid-β Peptides. Phys. Chem. Chem. Phys. 2017, 19 (24), 16257–16266. https://doi.org/10.1039/c6cp07774a.

(61) Terakawa, M. S.; Lin, Y.; Kinoshita, M.; Kanemura, S.; Itoh, D.; Sugiki, T.; Okumura, M.; Ramamoorthy, A.; Lee, Y. H. Impact of Membrane Curvature on Amyloid Aggregation. Biochim. Biophys. Acta - Biomembr. 2018, 1860 (9), 1741–1764. https://doi.org/10.1016/j.bbamem.2018.04.012.

(62) van Meer, G.; Voelker, D. R.; Feigenson, G. W. Membrane Lipids: Where They Are and How They Behave. Nat. Rev. Mol. Cell Biol. 2008, 9 (2), 112–124. https://doi.org/10.1038/nrm2330.

(63) Galvagnion, C.; Brown, J. W. P.; Ouberai, M. M.; Flagmeier, P.; Vendruscolo, M.; Buell, A. K.; Sparr, E.; Dobson, C. M. Chemical Properties of Lipids Strongly Affect the Kinetics of the Membrane-Induced Aggregation of α-Synuclein. Proc. Natl. Acad. Sci. 2016, 113 (26), 7065–7070. https://doi.org/10.1073/pnas.1601899113.

(64) Menon, S.; Sengupta, N.; Das, P. Nanoscale Interplay of Membrane Composition and Amyloid Self-Assembly. J. Phys. Chem. B 2020, 124 (28), 5837–5846. https://doi.org/10.1021/acs.jpcb.0c03796.

(65) Habchi, J.; Chia, S.; Galvagnion, C.; Michaels, T. C. T.; Bellaiche, M. M. J.; Ruggeri, F. S.; Sanguanini, M.; Idini, I.; Kumita, J. R.; Sparr, E.; Linse, S.; Dobson, C. M.; Knowles, T. P. J.; Vendruscolo, M. Cholesterol Catalyses Aβ42 Aggregation through a Heterogeneous Nucleation Pathway in the Presence of Lipid Membranes. Nat. Chem. 2018, 10 (6), 673– 683. https://doi.org/10.1038/s41557-018-0031-x.

(66) Dias, C. L.; Jalali, S.; Yang, Y.; Cruz, L. Role of Cholesterol on Binding of Amyloid Fibrils to Lipid Bilayers. J. Phys. Chem. B 2020, 124 (15), 3036–3042. https://doi.org/10.1021/acs.jpcb.0c00485.

(67) Tang, J.; Alsop, R. J.; Backholm, M.; Dies, H.; Shi, A.-C.; Rheinstädter, M. C. Amyloid-β 25–35 Peptides Aggregate into Cross-β Sheets in Unsaturated Anionic Lipid Membranes at High Peptide Concentrations. Soft Matter 2016, 12 (13), 3165–3176. https://doi.org/10.1039/C5SM02619A.

(68) Amaro, M.; Šachl, R.; Aydogan, G.; Mikhalyov, I. I.; Vácha, R.; Hof, M. GM1Ganglioside Inhibits β-Amyloid Oligomerization Induced by Sphingomyelin. Angew. Chemie - Int. Ed. 2016, 55 (32), 9411–9415. https://doi.org/10.1002/anie.201603178.

(69) Koppaka, V.; Axelsen, P. H. Accelerated Accumulation of Amyloid β Proteins on Oxidatively Damaged Lipid Membranes. Biochemistry 2000, 39 (32), 10011–10016. https://doi.org/10.1021/bi000619d.

(70) Zhang, X.; St Clair, J. R.; London, E.; Raleigh, D. P. Islet Amyloid Polypeptide Membrane Interactions: Effects of Membrane Composition. Biochemistry 2017, 56 (2), 376–390. https://doi.org/10.1021/acs.biochem.6b01016.

(71) Sciacca, M. F. M.; Brender, J. R.; Lee, D. K.; Ramamoorthy, A. Phosphatidylethanolamine Enhances Amyloid Fiber-Dependent Membrane Fragmentation. Biochemistry 2012, 51 (39), 7676–7684. https://doi.org/10.1021/bi3009888.

(72) Aisenbrey, C.; Borowik, T.; Byström, R.; Bokvist, M.; Lindström, F.; Misiak, H.; Sani, M. A.; Gröbner, G. How Is Protein Aggregation in Amyloidogenic Diseases Modulated by Biological Membranes? Eur. Biophys. J. 2008, 37 (3), 247–255. https://doi.org/10.1007/s00249-007-0237-0.

(73) Simons, K.; Vaz, W. L. C. Model Systems, Lipid Rafts, and Cell Membranes. Annu. Rev. Biophys. Biomol. Struct. 2004, 33 (1), 269–295. https://doi.org/10.1146/annurev.biophys.32.110601.141803.

(74) Shahane, G.; Ding, W.; Palaiokostas, M.; Orsi, M. Physical Properties of Model Biological Lipid Bilayers: Insights from All-Atom Molecular Dynamics Simulations. J. Mol. Model. 2019, 25 (3), 1–13. https://doi.org/10.1007/s00894-019-3964-0.

(75) John, T.; Thomas, T.; Abel, B.; Wood, B. R.; Chalmers, D. K.; Martin, L. L. How Kanamycin A Interacts with Bacterial and Mammalian Mimetic Membranes. Biochim. Biophys. Acta - Biomembr. 2017, 1859 (11), 2242–2252. https://doi.org/10.1016/j.bbamem.2017.08.016.

(76) Mattila, J.-P.; Sabatini, K.; Kinnunen, P. K. J. Oxidized Phospholipids as Potential Molecular Targets for Antimicrobial Peptides. Biochim. Biophys. Acta - Biomembr. 2008, 1778 (10), 2041–2050. https://doi.org/10.1016/j.bbamem.2008.03.020.

(77) Makky, A.; Tanaka, M. Impact of Lipid Oxidization on Biophysical Properties of Model Cell Membranes. J. Phys. Chem. B 2015, 119 (18), 5857–5863. https://doi.org/10.1021/jp512339m.

(78) Ferreira, T. M.; Sood, R.; Bärenwald, R.; Carlström, G.; Topgaard, D.; Saalwächter, K.; Kinnunen, P. K. J.; Ollila, O. H. S. Acyl Chain Disorder and Azelaoyl Orientation in Lipid Membranes Containing Oxidized Lipids. Langmuir 2016, 32 (25), 6524–6533. https://doi.org/10.1021/acs.langmuir.6b00788.

(79) Finkel, T.; Holbrook, N. J. Oxidants, Oxidative Stress and the Biology of Ageing. Nature 2000, 408 (6809), 239–247. https://doi.org/10.1038/35041687.

(80) Murray, I. V. J.; Liu, L.; Komatsu, H.; Uryu, K.; Xiao, G.; Lawson, J. A.; Axelsen, P. H. Membrane-Mediated Amyloidogenesis and the Promotion of Oxidative Lipid Damage by Amyloid Beta Proteins. J. Biol. Chem. 2007, 282 (13), 9335–9345. https://doi.org/10.1074/jbc.M608589200.

(81) Vus, K.; Girych, M.; Trusova, V.; Gorbenko, G.; Kinnunen, P.; Mizuguchi, C.; Saito, H. Fluorescence Study of the Effect of the Oxidized Phospholipids on Amyloid Fibril Formation by the Apolipoprotein A-I N-Terminal Fragment. Chem. Phys. Lett. 2017, 688, 1–6. https://doi.org/10.1016/j.cplett.2017.09.037.

(82) Pande, A. H.; Kar, S.; Tripathy, R. K. Oxidatively Modified Fatty Acyl Chain Determines Physicochemical Properties of Aggregates of Oxidized Phospholipids. Biochim. Biophys. Acta - Biomembr. 2010, 1798 (3), 442–452. https://doi.org/10.1016/j.bbamem.2009.12.028.

(83) Levine, H. Thioflavine T Interaction with Synthetic Alzheimer’s Disease B-amyloid Peptides: Detection of Amyloid Aggregation in Solution. Protein Sci. 1993, 2 (3), 404–410. https://doi.org/10.1002/pro.5560020312.

(84) Wolfe, L. S.; Calabrese, M. F.; Nath, A.; Blaho, D. V; Miranker, A. D.; Xiong, Y. Protein-Induced Photophysical Changes to the Amyloid Indicator Dye Thioflavin T. Proc. Natl. Acad. Sci. 2010, 107 (39), 16863–16868. https://doi.org/10.1073/pnas.1002867107.

(85) Mattila, J. P.; Sabatini, K.; Kinnunen, P. K. J. Oxidized Phospholipids as Potential Novel Drug Targets. Biophys. J. 2007, 93 (9), 3105–3112. https://doi.org/10.1529/biophysj.107.103887.

(86) Haberland, M. E.; Reynolds, J. A. Self Association of Cholesterol in Aqueous Solution. Proc. Natl. Acad. Sci. U. S. A. 1973, 70 (8), 2313–2316. https://doi.org/10.1073/pnas.70.8.2313.

(87) Avanti. Critical Micelle Concentrations (CMCs) https://avantilipids.com/tech-support/physical-properties/cmcs.

(88) Dahse, K.; Garvey, M.; Kovermann, M.; Vogel, A.; Balbach, J.; Fändrich, M.; Fahr, A. DHPC Strongly Affects the Structure and Oligomerization Propensity of Alzheimer’s Aβ(1- 40) Peptide. J. Mol. Biol. 2010, 403 (4), 643–659. https://doi.org/10.1016/j.jmb.2010.09.021.

(89) Sanders, H. M.; Jovcevski, B.; Marty, M. T.; Pukala, T. L. Structural and Mechanistic Insights into Amyloid-β and α-Synuclein Fibril Formation and Polyphenol Inhibitor Efficacy in Phospholipid Bilayers. FEBS J. 2021, 289, 215–230. https://doi.org/10.1111/febs.16122.

(90) John, T.; Gladytz, A.; Kubeil, C.; Martin, L. L.; Risselada, H. J.; Abel, B. Impact of Nanoparticles on Amyloid Peptide and Protein Aggregation: A Review with a Focus on Gold Nanoparticles. Nanoscale 2018, 10 (45), 20894–20913. https://doi.org/10.1039/C8NR04506B.

(91) John, T.; Adler, J.; Elsner, C.; Petzold, J.; Krueger, M.; Martin, L. L.; Huster, D.; Risselada, H. J.; Abel, B. Mechanistic Insights into the Size-Dependent Effects of Nanoparticles on Inhibiting and Accelerating Amyloid Fibril Formation. J. Colloid Interface Sci. 2022, 622, 804–818. https://doi.org/10.1016/j.jcis.2022.04.134.

(92) Grigolato, F.; Arosio, P. The Role of Surfaces on Amyloid Formation. Biophys. Chem. 2021, 270, 106533. https://doi.org/10.1016/j.bpc.2020.106533.

(93) Gladytz, A.; Abel, B.; Risselada, H. J. Gold-Induced Fibril Growth: The Mechanism of Surface-Facilitated Amyloid Aggregation. Angew. Chemie - Int. Ed. 2016, 55 (37), 11242– 11246. https://doi.org/10.1002/anie.201605151.

(94) Delgado, D. A.; Doherty, K.; Cheng, Q.; Kim, H.; Xu, D.; Dong, H.; Grewer, C.; Qiang, W. Distinct Membrane Disruption Pathways Are Induced by 40-Residue β-Amyloid Peptides. J. Biol. Chem. 2016, 291 (23), 12233–12244. https://doi.org/10.1074/jbc.M116.720656.

(95) Greenfield, N. J. Using Circular Dichroism Spectra to Estimate Protein Secondary Structure. Nat. Protoc. 2006, 1 (6), 2876–2890. https://doi.org/10.1038/nprot.2006.202.Using.

(96) Aisenbrey, C.; Bechinger, B.; Gröbner, G. Macromolecular Crowding at Membrane Interfaces: Adsorption and Alignment of Membrane Peptides. J. Mol. Biol. 2008, 375 (2), 376–385. https://doi.org/10.1016/j.jmb.2007.10.053.

(97) John, T.; Greene, G. W.; Patil, N. A.; Dealey, T. J. A.; Hossain, M. A.; Abel, B.; Martin, L.L. Adsorption of Amyloidogenic Peptides to Functionalized Surfaces Is Biased by Charge and Hydrophilicity. Langmuir 2019, 35 (45), 14522–14531. https://doi.org/10.1021/acs.langmuir.9b02063.

(98) Bokvist, M.; Lindström, F.; Watts, A.; Gröbner, G. Two Types of Alzheimer’s β-Amyloid (1-40) Peptide Membrane Interactions: Aggregation Preventing Transmembrane Anchoring Versus Accelerated Surface Fibril Formation. J. Mol. Biol. 2004, 335 (4), 1039–1049. https://doi.org/10.1016/j.jmb.2003.11.046.

(99) John, T.; Abel, B.; Martin, L. L. The Quartz Crystal Microbalance with Dissipation Monitoring (QCM-D) Technique Applied to the Study of Membrane-Active Peptides. Aust. J. Chem. 2018, 71 (7), 543–546. https://doi.org/10.1071/CH18129.

(100) Mechler, A.; Praporski, S.; Atmuri, K.; Boland, M.; Separovic, F.; Martin, L. L. Specific and Selective Peptide-Membrane Interactions Revealed Using Quartz Crystal Microbalance. Biophys. J. 2007, 93 (11), 3907–3916. https://doi.org/10.1529/biophysj.107.116525.

(101) McCubbin, G. A.; Praporski, S.; Piantavigna, S.; Knappe, D.; Hoffmann, R.; Bowie, J. H.; Separovic, F.; Martin, L. L. QCM-D Fingerprinting of Membrane-Active Peptides. Eur. Biophys. J. 2011, 40 (4), 437–446. https://doi.org/10.1007/s00249-010-0652-5.

(102) Janshoff; Galla; Steinem. Piezoelectric Mass-Sensing Devices as Biosensors-An Alternative to Optical Biosensors? Angew. Chem. Int. Ed. Engl. 2000, 39 (22), 4004–4032. https://doi.org/10.1002/1521-3773(20001117)39:22<4004::aid-anie4004>3.0.co;2-2.

(103) Hess, B.; Kutzner, C.; van der Spoel, D.; Lindahl, E. GROMACS 4: Algorithms for Highly Efficient, Load-Balanced, and Scalable Molecular Simulation. J. Chem. Theory Comput. 2008, 4 (3), 435–447. https://doi.org/10.1021/ct700301q.

(104) Ferreira, T. M.; Coreta-Gomes, F.; Ollila, O. H. S.; Moreno, M. J.; Vaz, W. L. C.; Topgaard, D. Cholesterol and POPC Segmental Order Parameters in Lipid Membranes: Solid State 1 H– 13 C NMR and MD Simulation Studies. Phys. Chem. Chem. Phys. 2013, 15 (6), 1976–1989. https://doi.org/10.1039/C2CP42738A.

(105) Khandelia, H.; Mouritsen, O. G. Lipid Gymnastics: Evidence of Complete Acyl Chain Reversal in Oxidized Phospholipids from Molecular Simulations. Biophys. J. 2009, 96 (7), 2734–2743. https://doi.org/10.1016/j.bpj.2009.01.007.

(106) Sani, M. A.; Separovic, F. How Membrane-Active Peptides Get into Lipid Membranes. Acc. Chem. Res. 2016, 49 (6), 1130–1138. https://doi.org/10.1021/acs.accounts.6b00074.

(107) Bücker, R.; Seuring, C.; Cazey, C.; Veith, K.; García-Alai, M.; Grünewald, K.; Landau, M. The Cryo-EM Structures of Two Amphibian Antimicrobial Cross-β Amyloid Fibrils. Nat. Commun. 2022, 13 (1), 4356. https://doi.org/10.1038/s41467-022-32039-z.

(108) Fatafta, H.; Khaled, M.; Owen, M. C.; Sayyed-Ahmad, A.; Strodel, B. Amyloid-β Peptide Dimers Undergo a Random Coil to β-Sheet Transition in the Aqueous Phase but Not at the Neuronal Membrane. Proc. Natl. Acad. Sci. 2021, 118 (39), e2106210118. https://doi.org/10.1073/pnas.2106210118.

(109) Humphrey, W.; Dalke, A.; Schulten, K. VMD: Visual Molecular Dynamics. J. Mol. Graph. 1996, 14 (1), 33–38. https://doi.org/10.1016/0263-7855(96)00018-5.

(110) Vácha, R.; Linse, S.; Lund, M. Surface Effects on Aggregation Kinetics of Amyloidogenic Peptides. J. Am. Chem. Soc. 2014, 136 (33), 11776–11782. https://doi.org/10.1021/ja505502e.

(111) Tsubone, T. M.; Junqueira, H. C.; Baptista, M. S.; Itri, R. Contrasting Roles of Oxidized Lipids in Modulating Membrane Microdomains. BBA - Biomembr. 2019, 1861 (3), 660– 669. https://doi.org/10.1016/j.bbamem.2018.12.017.

(112) Schumann-Gillett, A.; O’Mara, M. L. The Effects of Oxidised Phospholipids and Cholesterol on the Biophysical Properties of POPC Bilayers. Biochim. Biophys. Acta - Biomembr. 2019, 1861 (1), 210–219. https://doi.org/10.1016/j.bbamem.2018.07.012.

(113) Boonnoy, P.; Jarerattanachat, V.; Karttunen, M.; Wong-Ekkabut, J. Bilayer Deformation, Pores, and Micellation Induced by Oxidized Lipids. J. Phys. Chem. Lett. 2015, 6 (24), 4884–4888. https://doi.org/10.1021/acs.jpclett.5b02405.

(114) Wallgren, M.; Beranova, L.; Pham, Q. D.; Linh, K.; Lidman, M.; Procek, J.; Cyprych, K.; Kinnunen, P. K. J.; Hof, M.; Gröbner, G. Impact of Oxidized Phospholipids on the Structural and Dynamic Organization of Phospholipid Membranes: A Combined DSC and Solid State NMR Study. Faraday Discuss. 2012, 161, 499–513. https://doi.org/10.1039/c2fd20089a.

(115) Espinosa, Y. R.; Barrera Valderrama, D. I.; Carlevaro, C. M.; Llanos, E. J. Molecular Basis of the Anchoring and Stabilization of Human Islet Amyloid Polypeptide in Lipid Hydroperoxidized Bilayers. Biochim. Biophys. Acta - Gen. Subj. 2022, 1866 (10), 130200. https://doi.org/10.1016/j.bbagen.2022.130200.

(116) Yang, Y.; Jalali, S.; Nilsson, B. L.; Dias, C. L. Binding Mechanisms of Amyloid-like Peptides to Lipid Bilayers and Effects of Divalent Cations. ACS Chem. Neurosci. 2021, 12 (11), 2027–2035. https://doi.org/10.1021/acschemneuro.1c00140.

