## Supporting Information for "Lipid Oxidation Controls Peptide Self-Assembly near Membranes"

---

### 1 Materials and Methods

#### 1.1 Peptides and Buffer

Amyloid β (1−40) peptide (A$\beta_{40}$, DAEFRHDSGYEVHHQKLVFFAEDVGSNKGAIIGLMVGGVV, 95%) was obtained from Peptide 2.0 Inc. (Chantilly, VA) (used for all experiments and data shown) and Pepscan (Lelystad, Netherlands) (used for verification experiments). Uperin 3.5 peptide (U3.5, GVGDLIRKAVSVIKNIV-$NH_2$, 98%) was purchased from Peptide 2.0 Inc. (Chantilly, VA). Potassium phosphate monobasic (≥99.0%), potassium phosphate dibasic (≥98.0%) and sodium chloride (≥99.5%) were obtained from Sigma-Aldrich Corp. (St. Louis, MO). Ultrapure water (18.2 MΩ cm) was used for all experiments (Sartorius, Göttingen, Germany). Phosphate-buffered saline (PBS) solutions (20 mM phosphate and 100 mM sodium chloride) were used as buffer. The pH was adjusted to 7.40±0.05 using diluted sodium hydroxide (≥97.0%, pellets) or hydrochloric acid (32%). PBS solutions were filtered through hydrophobic polypropylene membrane filters prior to use (GH Polypro, 0.2 µm, PALL Life Sciences, USA).

#### 1.2 Liposome Preparation

1-Palmitoyl-2-oleoyl-*sn*-glycero-3-phosphocholine (POPC, >98%), 1-palmitoyl-2-oleoyl-*sn*-glycero-3-phospho-(1'-*rac*-glycerol) (POPG, >98%) and 1-palmitoyl-2-azelaoyl-sn-glycero-3-phosphocholine (PazePC, >98%) were obtained from Cayman Chemical (Ann Arbor, MI). Cholesterol (ovine wool, >98%) was obtained from Avanti Polar Lipids, Inc. (Alabaster, AL). POPC and POPG were each dissolved in a 2:1 (v/v) mixture of chloroform (≥99%, Sigma-Aldrich Corp., St. Louis, MO) and methanol (≥99.8%, Merck KGaA, Darmstadt, Germany), cholesterol was dissolved in pure ethanol-free chloroform and PazePC was diluted with methanol to prepare 5 mM lipid stock solutions. To mimic different membrane compositions, the following mixtures were prepared: POPC-POPG (4:1, molar ratio), POPC-cholesterol (4:1) and POPC-PazePC (7:3). Aliquots of 100 µL of each lipid or lipid mixture were transferred into test tubes, the solvent was evaporated under a gentle $N_2$ gas stream and dried under vacuum in a desiccator, sealed with parafilm and stored at -20°C. To form liposomes, the lipids were suspended in 0.5 mL of PBS buffer (1 mM lipid), incubated at 37°C for at least 30 min, vortexed for 5 min, and finally 15 times extruded through polycarbonate membranes (0.1 µm, Avanti Polar Lipids, Inc., Alabaster, AL). The lipids POPC, POPG and the mixtures POPC-POPG and POPC-Cholesterol were in liquid phase state under experimental conditions at 22°C or 37°C.[1]

#### 1.3 Dynamic Light Scattering (DLS) of Liposomes and Micelles

DLS measurements were performed to assess the hydrodynamic diameter (size) of liposomes using a Zetasizer Nano ZS instrument from Malvern Instruments (Malvern, UK). Liposome solutions were diluted (1:100) and measured in disposable solvent-resistant microcuvettes (ZEN0040, Malvern Instruments, Malvern, UK) at 25°C at a scattering angle of 173° (Non-Invasive Backscatter, NIBS). Sample material was set to protein (refractive index 1.450, absorption 0.001) due to its similar properties to lipids and dispersant was water (viscosity 0.8872 mPas, refractive index 1.330). Number density distributions are presented.

#### 1.4 Thioflavin T (ThT) Fluorescence Assays

ThT fluorescence assays were performed to study the kinetics of amyloid fibril formation, as previously reported.[2] ThT (Sigma-Aldrich Corp., St. Louis, MO) was diluted in dimethyl sulfoxide (DMSO, ≥99.9%, Merck, Darmstadt, Germany) to obtain a 1 mM ThT stock solution that was stored at -20 °C and protected from light. The peptides A$\beta_{40}$ (100 µM) and U3.5 (50 µM) were first dissolved in DMSO (20 µL of DMSO per each mL of the final sample volume) and ThT stock solution was added to obtain a ThT concentration of 20 µM in the final sample volume. Solutions of varying concentrations of lipid in PBS buffer were prepared and subsequently added to the peptide-ThT mixtures to obtain the final sample solutions (0, 100, 900 µM for A$\beta_{40}$ and 0, 50, 450 µM for U3.5) and initiate peptide aggregation kinetics. All sample solutions were vortexed before the transfer of 50 µL (for A$\beta_{40}$) or 150 µL (for U3.5) into each well of black polystyrene 96-well microplates with a solid, clear bottom, and nonbinding coating (Greiner Bio-One International GmbH, Kremsmünster, Austria). Microplates were closed with ThinSeal adhesive sealing (Astral Scientific Pty. Ltd., Taren Point, Australia) films to prevent evaporation. ThT fluorescence was recorded using a CLARIOstar plate reader (BMG LABTECH GmbH, Ortenberg, Germany) with excitation and emission wavelengths set to $\lambda_{exc}$ = 440 nm (bandwidth 10 nm) and $\lambda_{em}$ = 480 nm (bandwidth 10 nm), respectively. Experiments were performed at least in triplicate at 37°C, and the microplate was agitated for 40 seconds before each measurement cycle (5 minutes) using double orbital shaking (300 rpm). ThT data for A$\beta_{40}$ showed higher variations between repetitions. Data were averaged, normalized to a maximum fluorescence of 1 (except in cases with inhibition of peptide aggregation) and plotted in Origin 2022 (OriginLab Corp., Northampton, MA).

#### 1.5 Circular Dichroism (CD) Spectroscopy

A$\beta_{40}$ was first dissolved in a mixture of acetonitrile and 2,2,2-trifluoroethanol/TFE (1:1, v/v, 40 µL per each mL of final sample volume), and U3.5 was dissolved in a mixture of acetonitrile and ultrapure water (1:1, v/v, 40 µL per each mL of final sample volume) to prepare stock solutions. Acetonitrile (≥99.9%) was obtained from Merck (Darmstadt, Germany) and 2,2,2-trifluoroethanol (TFE, ≥99.0%) was obtained from Sigma-Aldrich Corp. (St. Louis, MO). Solvents were adjusted, different to the ThT assays, to consider that DMSO would absorb in the region of interest for CD measurements. The peptide stock solutions were aliquoted and liposomes in PBS buffer were added to obtain final peptide concentrations of 100 µM ($A\beta_{40}$) or 50 µM (U3.5) and liposome concentrations of 900 µM ($A\beta_{40}$) or 450 µM (U3.5). Samples were measured by CD spectroscopy immediately after sample preparation and after 15 h (U3.5) or two days ($A\beta_{40}$) of incubation at 37°C. U3.5 samples were directly measured, whereas aliquots of the $A\beta_{40}$ samples were diluted to 20 µM peptide / 180 µM liposomes due to a higher CD absorbance.

Samples were vortexed before the transfer of 100 µL ($A\beta_{40}$) or 150 µL (U3.5) sample solution into 1 mm path length quartz cuvettes (21/10/Q/1/ CD, Starna Scientific Ltd., Essex, UK). Samples were mixed before each measurement by inverting the cuvettes multiple times for 30 s. Far-UV CD spectra were recorded between 260 and 195 nm at 37°C using a J-815 CD spectropolarimeter (Jasco Corp., Tokyo, Japan) at standard sensitivity as previously reported (DIT, 1 s; bandwidth, 1 nm; data pitch, 0.5 nm; continuous; scanning speed, 50 nm/min; five scans).[2] The buffer contribution was subtracted for each experiment, and experiments were repeated at least in duplicate. Representative data were plotted in Origin 2022 (OriginLab Corp., Northampton, MA). The mean residue molar ellipticities (MRE, $\Theta_{\text{molar},\lambda}$) were determined from the measured ellipticities $\Theta_\lambda$, the peptide concentrations $c$, the cuvette path lengths $l$ and the number of residues $n$, i.e., 39 peptide bonds for $A\beta_{40}$ and 16 peptide bonds for U3.5).

$$\theta_{\text{molar},\lambda} = \frac{\theta_\lambda \text{ (millidegrees) } 10^6}{c \text{ (micromolar) } l \text{ (millimeters) } n}$$

The secondary structure content for the peptides was calculated from the measured CD spectra using the BeStSel webserver (https://bestsel.elte.hu) ($\alpha$-helix, β-strand, turns, and other).[3]

**1.6 Quartz Crystal Microbalance (QCM)**

Silicon dioxide (SiO2)-coated quartz crystals with a fundamental frequency of 5 MHz (Q-Sense, Biolin Scientific, Gothenburg, Sweden) were used as sensors for the QCM experiments. The sensors were cleaned using the following protocol: After 10 min in a 2% Hellmanex II cleaning solution (Hellma, Mülheim, Germany), the sensors were rinsed with water, followed by isopropanol (>99.5%, Merck, Germany) and dried under a gentle nitrogen gas stream. The sensors were finally treated in a UV Ozone ProCleaner (BioForce Nanosciences, Virginia Beach, VA) for 20 min.

QCM measurements were performed using a Q-Sense E4 instrument (Biolin Scientific, Gothenburg, Sweden) consisting of four flow cells at 22±0.05 °C at least in triplicate. Initially, ultrapure water, followed by PBS buffer, were introduced into the measurement flow cells at 200 μL/min to obtain baseline signals. Liposomes (0.1 mM lipid in 20 mM phosphate and 100 mM (for POPC) / 250 mM (for POPG) sodium chloride buffer) were deposited onto the silicon dioxide sensors at a flow rate of 50 μL/min until they ruptured and formed a lipid bilayer structure. This was visible from a sudden increase in frequency, followed by stable frequency values.[4,5] An increase in frequency ($\Delta f$) is correlated with a decrease in mass ($\Delta m$) through the Sauerbrey equation ($\Delta f = - C \cdot \Delta m$).[6] The measurement cells were rinsed with phosphate buffer at 200 μL/min to remove any lipids that were not bound to the sensors, followed by the introduction of peptide sample solution at 25 μM at a flow rate of 50 μL/min for 15 min. Frequency changes were followed for another 45 min without flow under steady conditions, followed by a final rinse with PBS buffer for at least 10 min.

Changes in frequency ($\Delta f$) were measured over time while the quartz crystal sensors were excited at multiple harmonics (n = 1, 3, 5, 7, 9 and 11; respectively 5–55 MHz). Data for the 7$^{th}$ harmonic were used. All frequency values ($\Delta f$) were normalized to the fundamental frequency ($\Delta f_n/n$). QCM data were exported from QTools (Q-Sense, Biolin Scientific, Gothenburg, Sweden) and analyzed in Origin 2022 (OriginLab Corporation, Northampton, MA).

**1.7 Molecular Dynamics (MD) Simulations**

MD simulations were performed at 303.15 K using the GROMACS 4.5.7 software package.[7–10] Model membranes were simulated with lipid compositions consistent with experimental conditions to understand peptide-membrane interactions at the molecular detail. The GROMOS 54A7 united-atom force field was used to describe the peptides, water and ions.[11] All lipid force fields were used as previously reported in the literature.[12,13] POPC molecules were parametrized based on the Berger force field[14] with double bond correction introduced by Bachar *et al.*[15,16] The cholesterol force field is GROMOS based.[17] The molecule types $CH_2$/$CH_3$ in the cholesterol force field were changed to avoid overcondensation of the bilayer as previously implemented.[12,18] The force field for deprotonated PazePC was obtained from Khandelia and Mouritsen[19] who derived it from the POPC force field by shortening the oleoyl chain at the double bond and replacing it with a carboxyl group, correcting geometry and using partial charges from amino acids. The force field for protonated PazePC was adapted from Khandelia and Mouritsen[19] by adding hydrogen to the deprotonated PazePC together with ad hoc partial charges to obtain a total charge of zero, as reported by Ferreira *et al.*[13] POPG was described using a Berger based force field.[20] The force field parameters for POPC, cholesterol and PazePC were received from Ollila *et al.*[21–24] and for POPG from the MemBuilder II webserver.[25]

Each lipid bilayer consisted of 128 lipid molecules, i.e. two layers of each 8x8 (64) lipid molecules. The structure files of the pure POPC bilayer and the POPC-PazePC bilayer (7:3; 90 POPC and 38 PazePC molecules) were obtained from Ollila *et al.*[21,23,24] The POPC-cholesterol bilayer structure (4:1; 102 POPC and 26 cholesterol molecules) was obtained from Ollila *et al.*[22] and manually adapted for the correct amount of cholesterol (8 POPC molecules were removed and manually replaced by cholesterol molecules). The POPC-POPG bilayer structure (4:1; 102 POPC and 26 POPG molecules) was generated using MemBuilder II.[25] The $A\beta_{40}$ peptide was used with both a random coil (PDB Utilities Server)[26] and an $\alpha$-helical (adapted from PDB: 1IYT)[27] starting structure from the Protein Data Bank (PDB), while the U3.5 peptide was studied with a random coil and an $\alpha$-helical starting structure, as obtained in nuclear magnetic resonance (NMR) experiments in sodium dodecyl sulfate (SDS) micelles.[28] The partially folded $3_{10}$ helix structure of $A\beta_{40}$ (PDB: 2LFM) could have been used alternatively.[29] N- and C-termini of the $A\beta_{40}$ peptide were charged, whereas the C-terminus of U3.5 was uncharged (amide modified).

The following simulation parameters were used for all simulations: Periodic boundary conditions were applied. The Particle Mesh Ewald (PME) method with a grid of 0.12 nm, a fourth order spline interpolation and a Coulomb cut-off at 1.0 nm was used to describe electrostatic interactions[30,31] and a Lennard-Jones cut-off distance of 1.0 nm was used to describe van der Waals interactions. The neighborlist was updated every tenth step with a time step of 2 fs. Centre of mass motion was removed for the system at every step. All bonds were constrained to their equilibrium values using the LINCS algorithm.[32] Explicit water (Simple Point Charge, SPC)[33] was constrained using the SETTLE algorithm.[34] The temperature was coupled separately for lipids and peptide/water/ions to 303.13 K using the velocity-rescale method with a coupling constant of 0.1 $ps^{-1}$.[35] The pressure was semiisotropically coupled to 1 bar with the Berendsen barostat.[36]

Prior to adding peptide, all lipid bilayers were equilibrated for at least 50 ns. One peptide molecule was randomly positioned outside the lipid bilayer and solvated with about 6000-7000 water molecules. 150 mM NaCl was added as salt and to electroneutralize the systems (see Figure S1 and Table S1 for an overview). All simulations were run for 100 ns in triplicate. Further, each peptide was simulated in water without any lipid present for 100 ns in triplicate, serving as a reference.

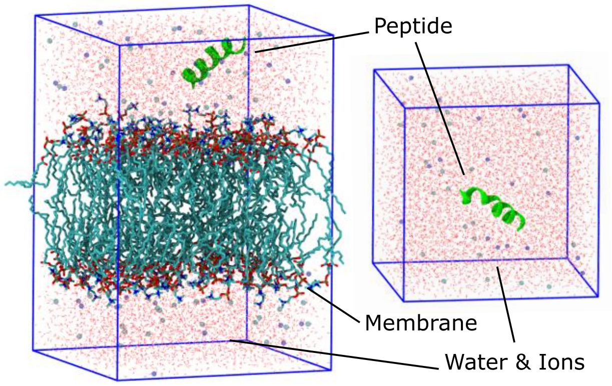


**Figure S1.** Representative initial simulation setups for the peptides with lipid bilayer (left) and without lipids (right) in solution. Snapshots of the simulation boxes with U3.5 peptide in helical secondary structure near a POPC membrane and in water are shown.

**Table S1.** Overview of MD Simulations (each 3 x 100 ns).

| Lipids | Peptides |
|---|---|
| **Bilayer Structures (128 lipid molecules)** | |
| POPC | 1 x A$\beta_{40}$ ($\alpha$-helix), 1 x A$\beta_{40}$ (random), 1 x U3.5 ($\alpha$-helix), 1 x U3.5 (random) |
| POPC-cholesterol (4:1) | 1 x A$\beta_{40}$ ($\alpha$-helix), 1 x A$\beta_{40}$ (random), 1 x U3.5 ($\alpha$-helix), 1 x U3.5 (random) |
| POPC-POPG (4:1) | 1 x A$\beta_{40}$ ($\alpha$-helix), 1 x A$\beta_{40}$ (random), 1 x U3.5 ($\alpha$-helix), 1 x U3.5 (random) |
| POPC-PazePC (protonated) (7:3) | 1 x A$\beta_{40}$ ($\alpha$-helix), 1 x A$\beta_{40}$ (random), 1 x U3.5 ($\alpha$-helix), 1 x U3.5 (random) |
| POPC-PazePC (deprotonated) (7:3) | 1 x A$\beta_{40}$ ($\alpha$-helix), 1 x A$\beta_{40}$ (random), 1 x U3.5 ($\alpha$-helix), 1 x U3.5 (random) |

MD simulation snapshots were visualized in VMD 1.93.[37] Representative structures of the simulation trajectories were determined using clustering analysis of the last 10 ns of all repetitions each (3 x 10 ns, gromos method, RMSD cutoff 0.2 nm, g_cluster).[38] The central structure of the largest clusters were visualized. The secondary structure content of the peptides was analyzed using the DSSP tool (Define Secondary Structure of Proteins, do_dssp) for the last 10 ns of simulation time.[39,40] Mass density profiles of the lipids, peptides, and water and ions were analyzed (g_density) for the last 10 ns simulation time and plotted perpendicular to the membrane plane. Minimum distances between the lipid membranes and each peptide residue were determined for the last 10 ns simulation time (g_pairdist). Average distances in $z$-dimension, perpendicular to the membranes, between phosphate atoms in POPC and peptide $C_\alpha$ atoms were calculated for the last 10 ns simulation time (g_traj). Averaged data of the repetitions were plotted in Origin 2022 (OriginLab Corporation, Northampton, MA). The lipid structures in the manuscript were prepared in ChemDraw 18.0 (PerkinElmer, Waltham, MA). Peptide properties were calculated using pepcalc.com (Innovagen AB, 2015).

### 2 Supporting Results

#### 2.1 Dynamic Light Scattering (DLS) of Liposomes and Micelles

In this work, the impact of different membrane components on peptide aggregation was studied. For most of the phospholipids studied, stable liposomes were formed, even at the lowest concentrations (see Figure S2). Pure PazePC has a critical micelle concentration (cmc) of about 20 µM.[41,42] The cmc for POPC, POPG and cholesterol (25-40 nM)[43] are in the nM range and thus micelles and liposomes are formed under all conditions used in this study (50-900 µM).[44]

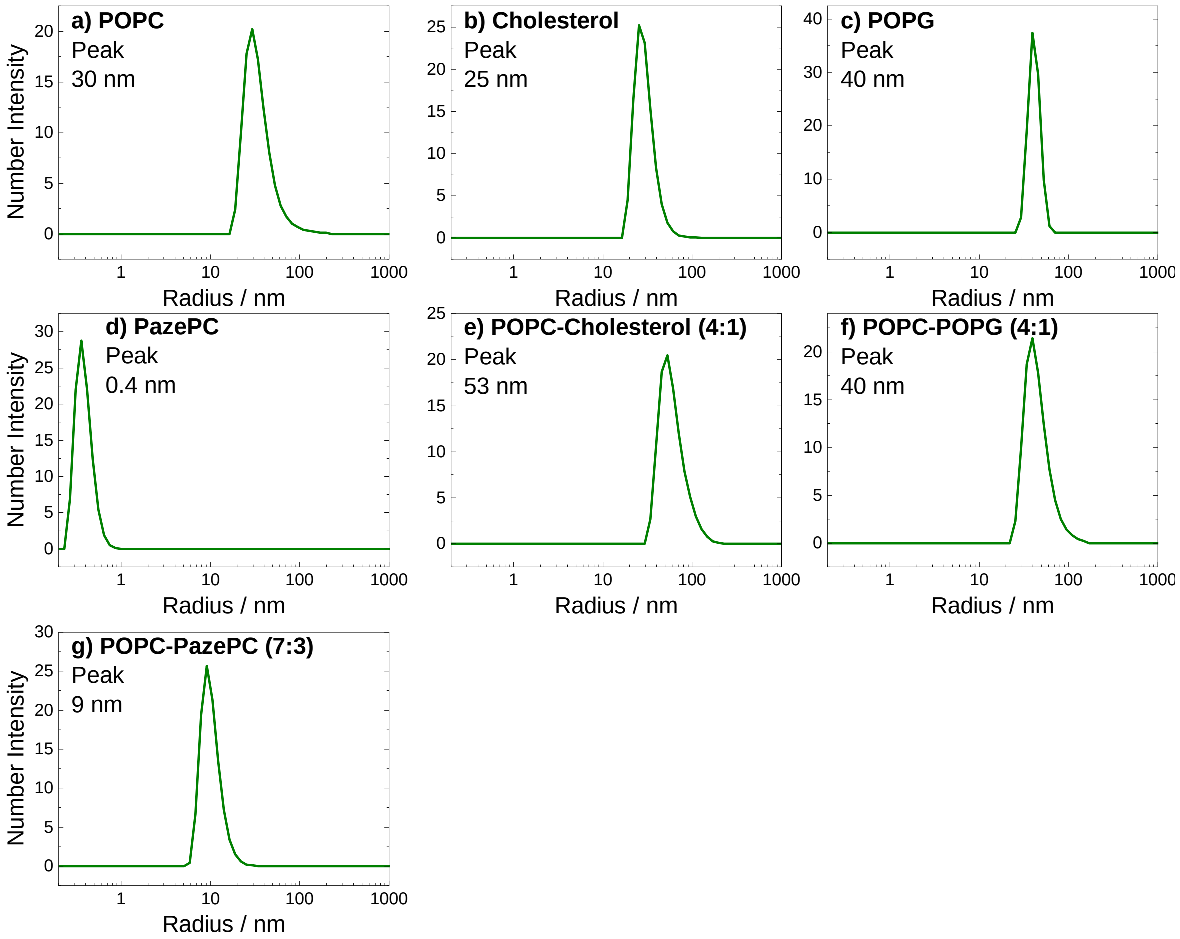


**Figure S2.** Hydrodynamic radii of lipids in aqueous PBS buffered solution at 25°C. DLS number intensity distributions are shown. Small unilamellar vesicles (SUVs) were formed for a) POPC, b) Cholesterol, c) POPG, e) POPC-Cholesterol (4:1, molar ratio), and f) POPC-POPG (4:1). In contrast, g) POPC-PazePC (7:3) self-assembled into micelles and d) pure PazePC did not self-assemble into any larger structures at low concentration (10 µM).

### 2.2 Thioflavin T (ThT) Fluorescence Assays

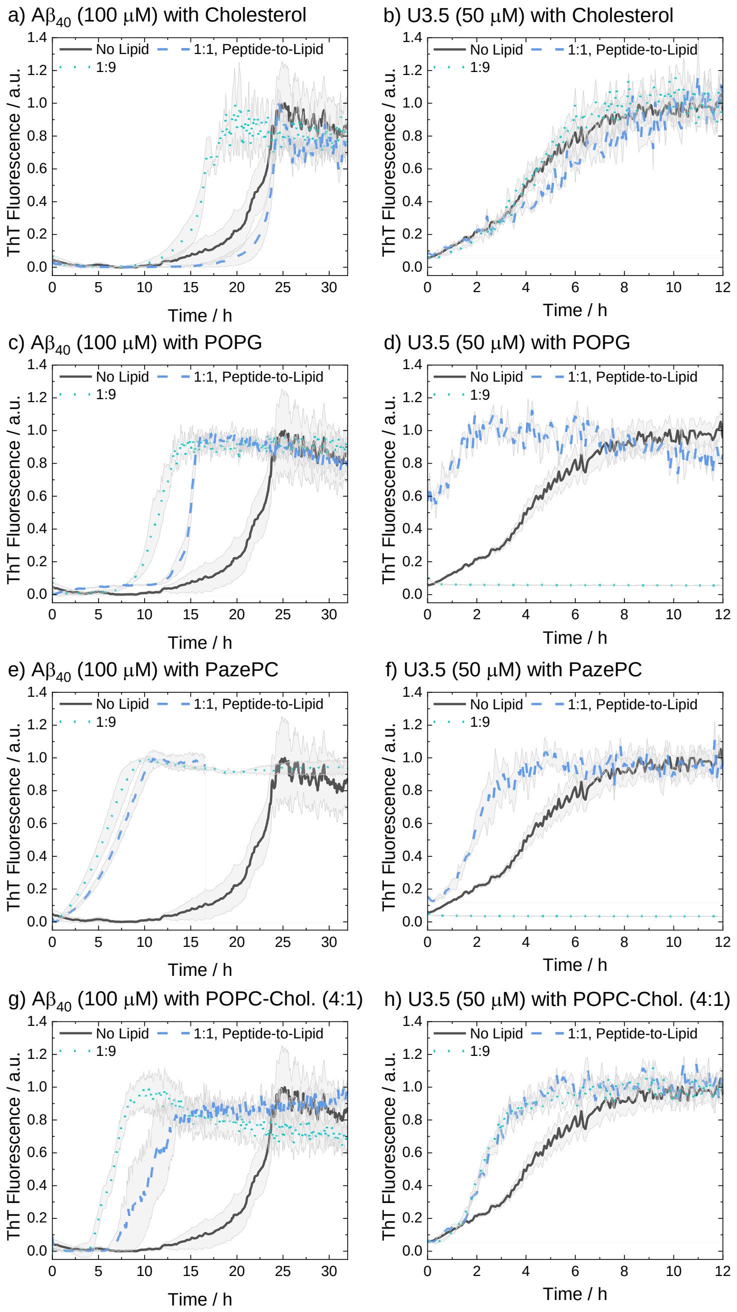


**Figure S3.** ThT fluorescence assays were performed to follow the kinetics of fibril formation. The peptides A$\beta_{40}$ (100 μM) and U3.5 (50 μM) were studied in PBS buffer at pH 7.4 at 37°C. Peptides were studied without and with different amounts of lipids present (peptide-to-lipid molar ratio: 1:1, 1:9). The largest impact of the lipids on peptide aggregation was observed when lipid was added in excess (1:9). When peptide and lipid had the same concentration in the sample (1:1), smaller effects were observed. Data for pure Cholesterol, POPG, and PazePC, and a mixture consisting of POPC-Cholesterol (4:1) are shown. The data for U3.5 (b) without lipid present and with excess of cholesterol were previously reported and are included as a reference to all other lipids and A$\beta_{40}$.[2] Data were normalized to a maximum fluorescence of 1 (except in cases with inhibition of peptide aggregation).

### 2.3 Circular Dichroism (CD) Spectroscopy

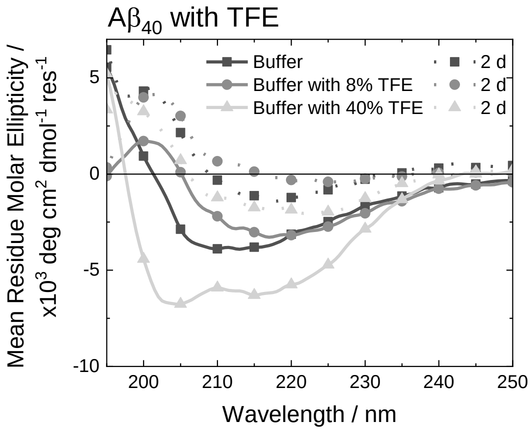


**Figure S4.** CD spectra of A$\beta_{40}$ without and with 2,2,2-trifluoroethanol (TFE) in PBS buffer at pH 7.4 at 37°C. A$\beta_{40}$ aggregation was studied at 100 μM with 8 % and 40 % of TFE present. For CD measurements, the samples were diluted to 20 μM peptide. Samples were measured initially (solid lines) and after 2 days (dotted lines). Note that the symbols are used to distinguish the data sets and data were recorded every 0.5 nm.

The secondary structure content for the peptides was calculated from the measured CD spectra using the BeStSel webserver (https://bestsel.elte.hu) ($\alpha$-helix, β-strand, turn, other) (Table S2).[3]

**Table S2.** Quantitative secondary structure estimation for the peptides A$\beta_{40}$ and U3.5 using BeStSel.[3] A low NRMSD value (normalized root mean square deviation) indicates a good fit. Secondary structures: $\alpha$-helix, β-strand, turn, other ($3_{10}$-helix, π-helix, bends, β-bridge, irregular/loop).

| Peptide and Condition | $\alpha$-helix | β-strand | Turns | Other | NRMSD |
|---|---|---|---|---|---|
| A$\beta_{40}$ in buffer after 2 days | 3.7 | 41.2 | 14.6 | 40.4 | 0.0312 |
| A$\beta_{40}$ with POPC after 2 days | 9.4 | 41.0 | 10.3 | 39.4 | 0.0072 |
| A$\beta_{40}$ with Cholesterol after 2 days | 4.6 | 32.3 | 14.4 | 48.6 | 0.0376 |
| A$\beta_{40}$ with POPG after 2 days | 4.1 | 32.2 | 14.8 | 49.0 | 0.0314 |
| A$\beta_{40}$ with PazePC after 2 days | 4.4 | 43.5 | 11.6 | 40.5 | 0.0094 |
| A$\beta_{40}$ with 8 % TFE | 3.0 | 36.1 | 13.6 | 47.2 | 0.0238 |
| A$\beta_{40}$ with 8 % TFE after 2 days | 3.0 | 42.0 | 14.5 | 40.5 | 0.0345 |
| A$\beta_{40}$ with 40 % TFE | 4.4 | 28.5 | 15.5 | 51.5 | 0.0165 |
| A$\beta_{40}$ with 40 % TFE after 2 days | 3.8 | 38.7 | 15.4 | 42.1 | 0.0216 |
| U3.5 in buffer after 15 hours | 3.0 | 23.4 | 11.8 | 61.8 | 0.0138 |
| U3.5 with POPC after 15 hours | 18.6 | 11.6 | 13.2 | 56.6 | 0.0114 |
| U3.5 with Cholesterol after 15 hours | 5.4 | 23.0 | 12.1 | 59.6 | 0.0121 |
| U3.5 with POPG after 15 hours | 55.9 | 8.9 | 9.8 | 25.5 | 0.0051 |
| U3.5 with PazePC after 15 hours | 56.2 | 1.8 | 9.6 | 32.4 | 0.0064 |

### 2.4 Molecular Dynamics (MD) Simulations

MD simulations were performed for the Aβ40 and U3.5 peptide with $\alpha$-helical and unstructured (random) starting structure each. While representative data are shown in the main manuscript, a comprehensive overview of the results of all simulations is included as part of the Supporting Information (Figures S5 – S12).

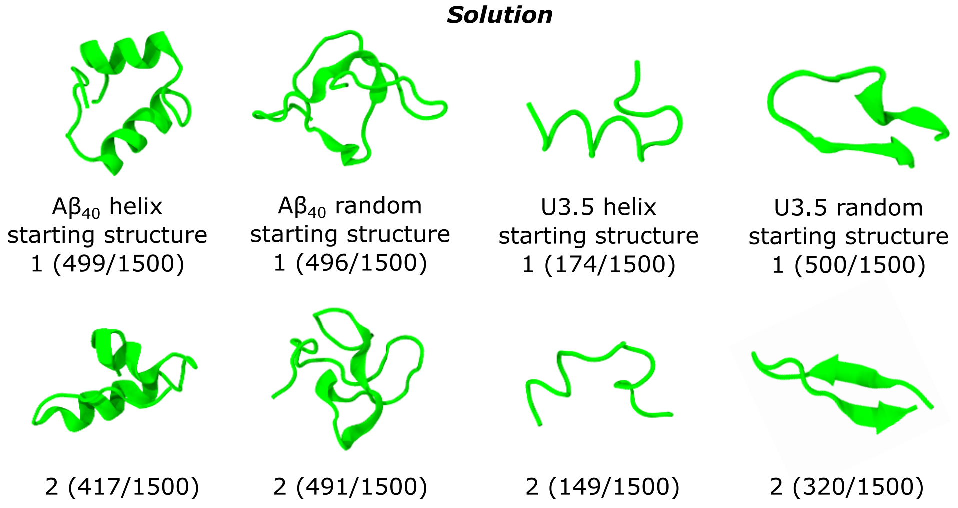


**Figure S5.** Cartoon representation of the peptides in solution, without lipids present. The central structure of the two largest structural clusters during the last 10 ns simulation time of all replicates for each peptide are shown. The numbers in parenthesis represent the size of the clusters of similar structures out of the 1500 frames each.

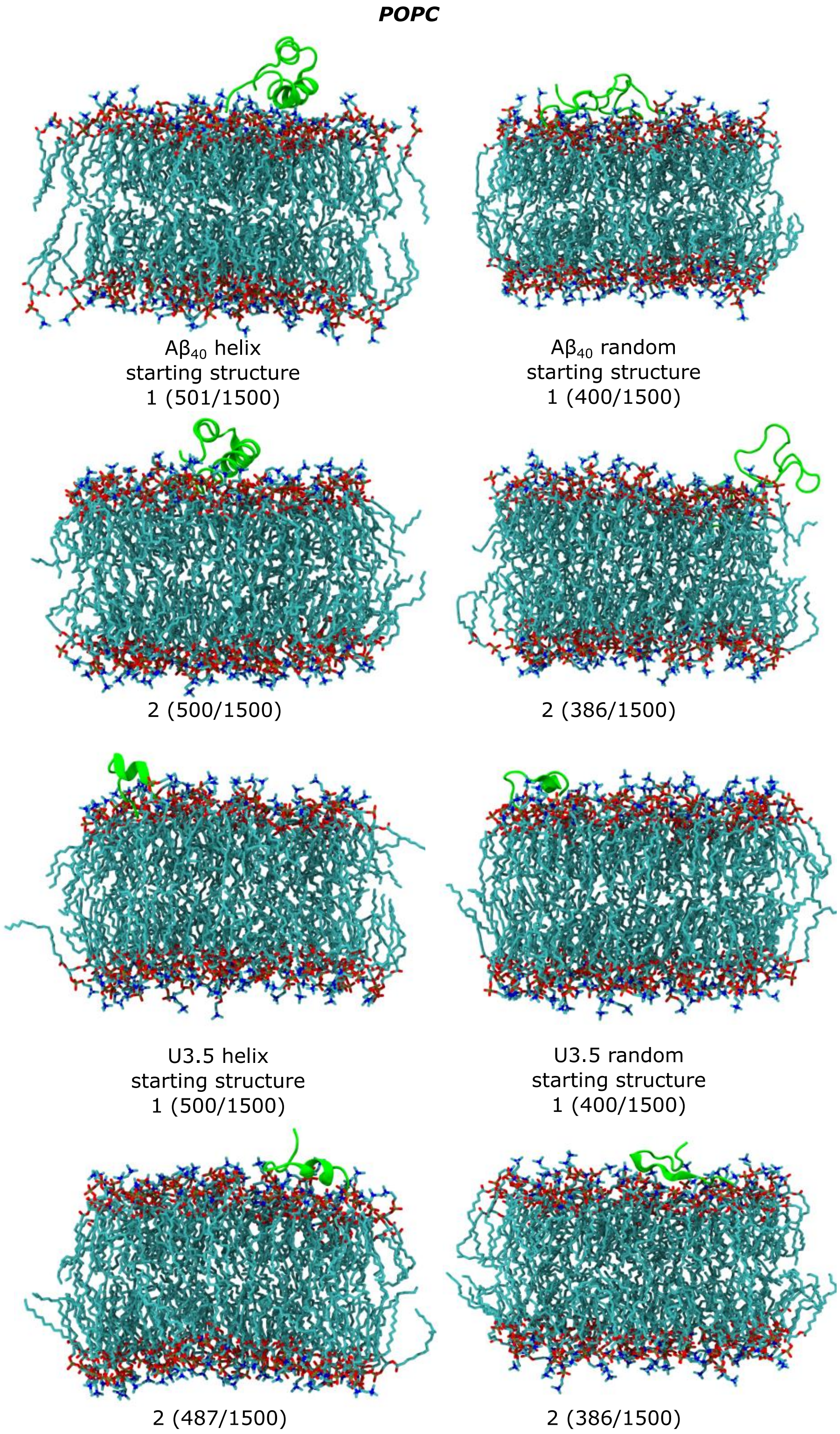


**Figure S6.** Cartoon representation of the peptides near POPC membranes. The central structure of the two largest structural clusters during the last 10 ns simulation time of all replicates for each peptide are shown. The numbers in parenthesis represent the size of the clusters of similar structures out of the 1500 frames each.

***POPC-Cholesterol (4:1)***

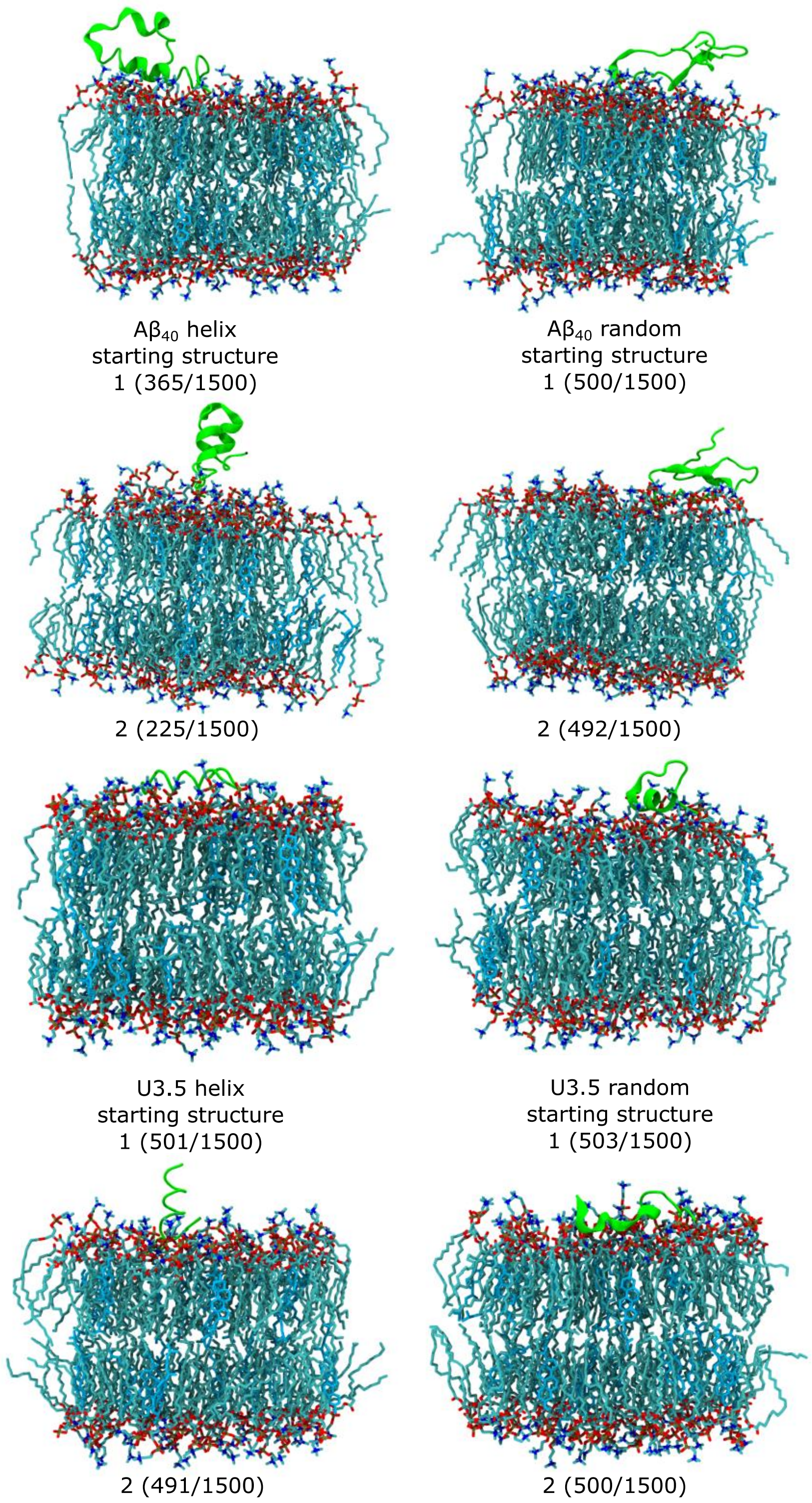


**Figure S7.** Cartoon representation of the peptides near POPC-Cholesterol (4:1) membranes. The central structure of the two largest structural clusters during the last 10 ns simulation time of all replicates for each peptide are shown. The numbers in parenthesis represent the size of the clusters of similar structures out of the 1500 frames each.

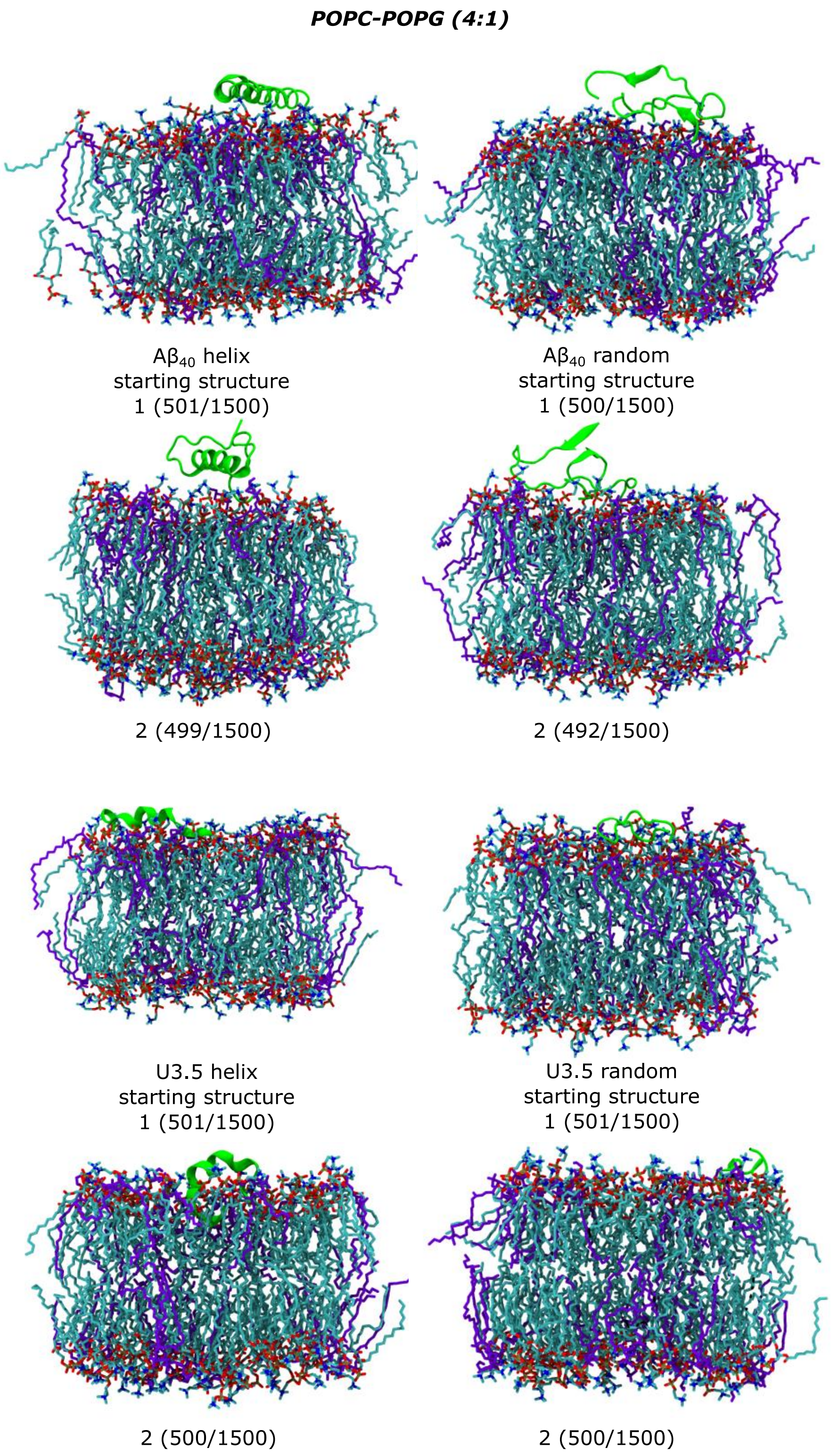


**Figure S8.** Cartoon representation of the peptides near POPC-POPG (4:1) membranes. The central structure of the two largest structural clusters during the last 10 ns simulation time of all replicates for each peptide are shown. The numbers in parenthesis represent the size of the clusters of similar structures out of the 1500 frames each.

***POPC-PazePC (protonated) (7:3)***

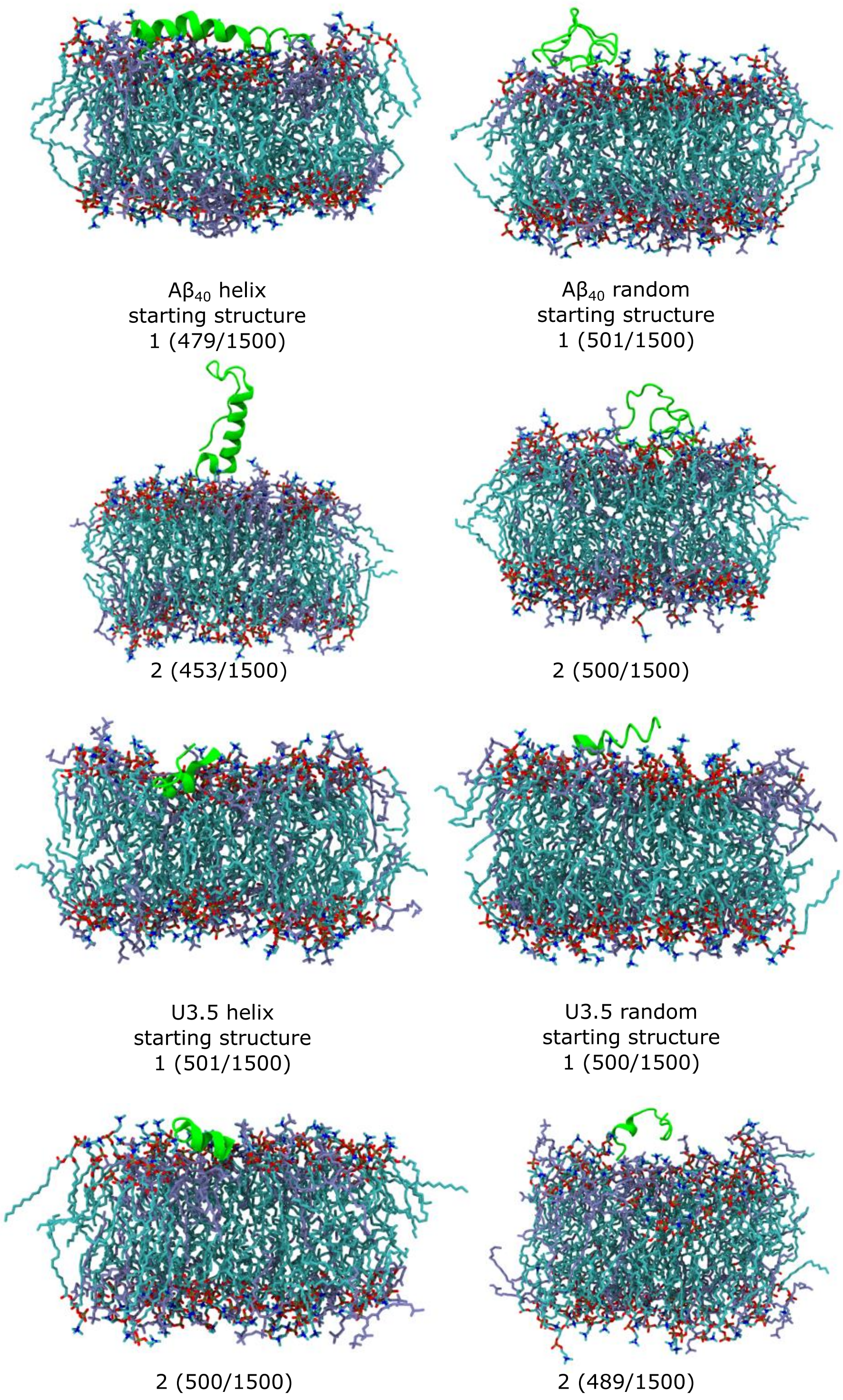


**Figure S9.** Cartoon representation of the peptides near POPC-PazePC (protonated) (7:3) membranes. The central structure of the two largest structural clusters during the last 10 ns simulation time of all replicates for each peptide are shown. The numbers in parenthesis represent the size of the clusters of similar structures out of the 1500 frames each.

***POPC-PazePC (deprotonated) (7:3)***

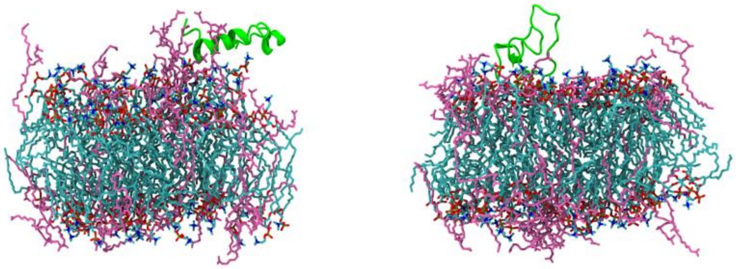

Aβ$_{40}$ helix starting structure 1 (217/1500)

Aβ$_{40}$ random starting structure 1 (501/1500)

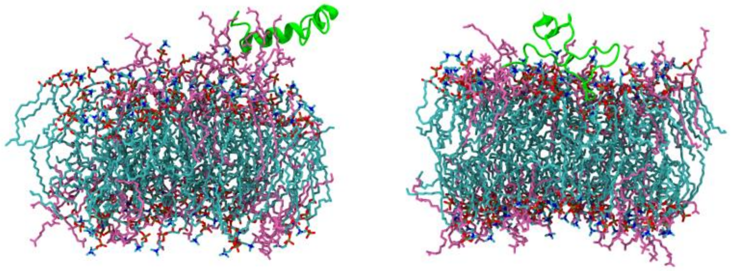

2 (212/1500)

2 (486/1500)

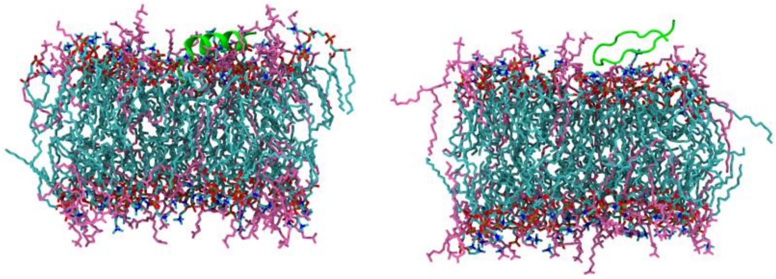

U3.5 helix starting structure 1 (501/1500)

U3.5 random starting structure 1 (501/1500)

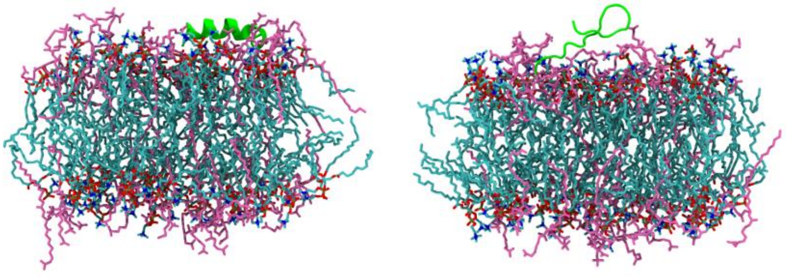

2 (499/1500)

2 (500/1500)

**Figure S10.** Cartoon representation of the peptides near POPC-PazePC (deprotonated) (7:3) membranes. The central structure of the two largest structural clusters during the last 10 ns simulation time of all replicates for each peptide are shown. The numbers in parenthesis represent the size of the clusters of similar structures out of the 1500 frames each.

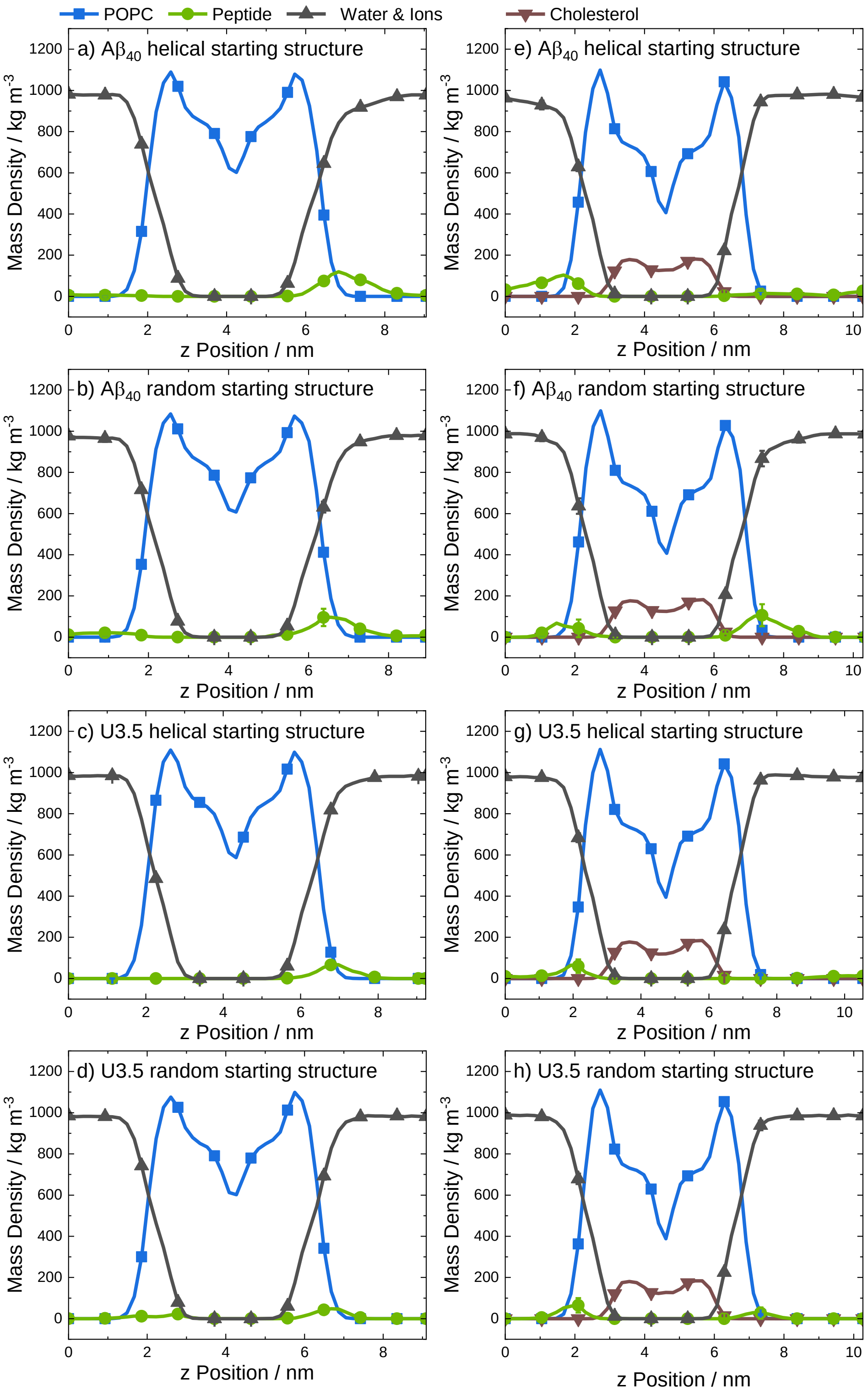


**Figure S11.** Averaged mass density profiles for the last 10 ns of all replicates for the simulations with POPC (left) and POPC-Cholesterol (4:1) (right) membranes. Coloring: POPC (blue), protein (green), water & ions (black), and cholesterol (brown). Note that the symbols are used to distinguish the data sets for every 5th data point for clarity.

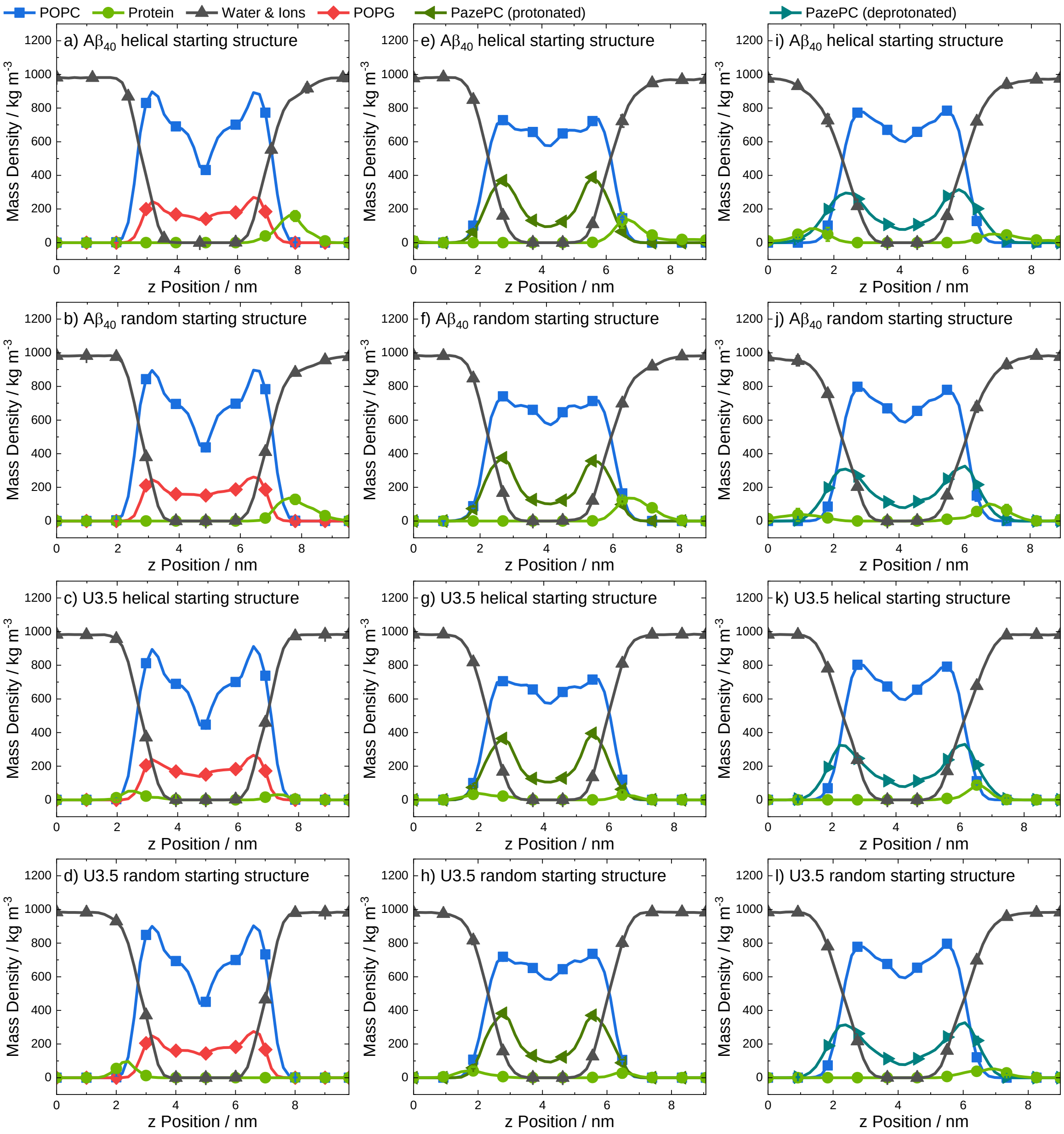


**Figure S12.** Averaged mass density profiles for the last 10 ns of all replicates for the simulations with POPC-POPG (4:1) (left), POPC-PazePC (protonated) (7:3) (center), and POPC-PazePC (deprotonated) (7:3) (right) membranes. Coloring: POPC (blue), protein (green), water & ions (black), POPG (red), PazePC (protonated) (dark green), and PazePC (deprotonated) (cyan). Note that the symbols are used to distinguish the data sets for every 5th data point for clarity.

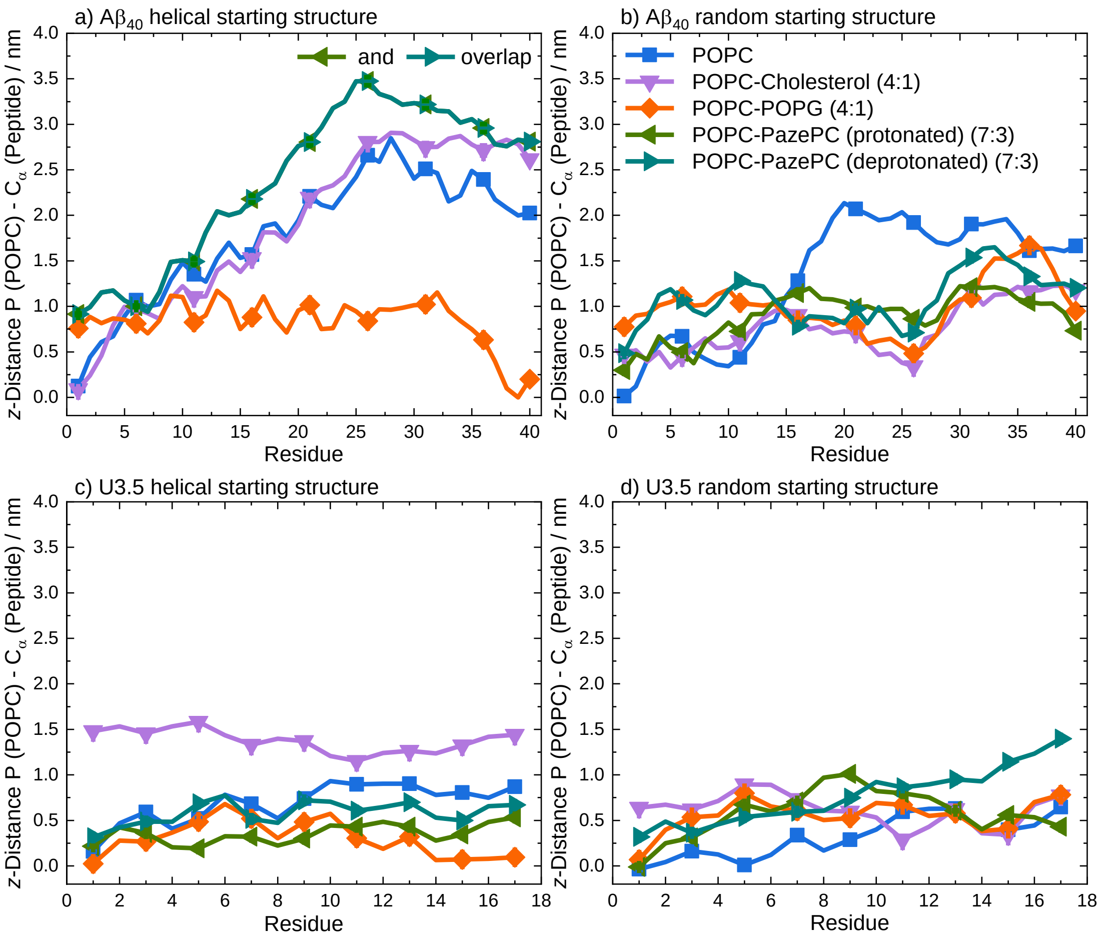


**Figure S13.** Average distances between the phosphate head groups of POPC in the outer membrane leaflet and the peptide Cα atoms of (a, b) Aβ40 and (c, d) U3.5 (perpendicular to the membrane along *z*-axis) during the last 10 ns simulation time of all replicates. U3.5 peptide shows stronger binding to the membrane surface compared to Aβ40. Note that the symbols are used to distinguish the data sets and each residue has a data point.

**3 Supporting References**


(1) Avanti. Phase Transition Temperatures for Glycerophospholipids https://avantilipids.com/tech-support/physical-properties/phase-transition-temps.

(2) John, T.; Dealey, T. J. A.; Gray, N. P.; Patil, N. A.; Hossain, M. A.; Abel, B.; Carver, J. A.; Hong, Y.; Martin, L. L. The Kinetics of Amyloid Fibrillar Aggregation of Uperin 3.5 Is Directed by the Peptide's Secondary Structure. *Biochemistry* **2019**, *58* (35), 3656–3668. https://doi.org/10.1021/acs.biochem.9b00536.

(3) Micsonai, A.; Wien, F.; Bulyáki, É.; Kun, J.; Moussong, É.; Lee, Y. H.; Goto, Y.; Réfrégiers, M.; Kardos, J. BeStSel: A Web Server for Accurate Protein Secondary Structure Prediction and Fold Recognition from the Circular Dichroism Spectra. *Nucleic Acids Res.* **2018**, *46* (W1), W315–W322. https://doi.org/10.1093/nar/gky497.

(4) Keller, C. A.; Kasemo, B. Surface Specific Kinetics of Lipid Vesicle Adsorption Measured with a Quartz Crystal Microbalance. *Biophys. J.* **1998**, *75* (3), 1397–1402. https://doi.org/10.1016/S0006-3495(98)74057-3.

(5) Cho, N.; Frank, C.; Kasemo, B.; Höök, F. Quartz Crystal Microbalance with Dissipation Monitoring of Supported Lipid Bilayers on Various Substrates. *Nat. Protoc.* **2010**, *5* (6), 1096–1106. https://doi.org/10.1038/nprot.2010.65.

(6) Sauerbrey, G. Verwendung von Schwingquarzen Zur Wägung Dünner Schichten Und Zur Mikrowägung. *Zeitschrift für Phys.* **1959**, *155* (2), 206–222. https://doi.org/10.1007/BF01337937.

(7) Hess, B.; Kutzner, C.; van der Spoel, D.; Lindahl, E. GROMACS 4: Algorithms for Highly Efficient, Load-Balanced, and Scalable Molecular Simulation. *J. Chem. Theory Comput.* **2008**, *4*, 435–447.

(8) van der Spoel, D.; Lindahl, E.; Hess, B.; Groenhof, G.; Mark, A. E.; Berendsen, H. J. C. GROMACS: Fast, Flexible, and Free. *J. Comput. Chem.* **2005**, *26*, 1701–1718.

(9) Lindahl, E.; Hess, B.; van der Spoel, D. GROMACS 3.0: A Package for Molecular Simulation and Trajectory Analysis. *J. Mol. Model.* **2001**, *7*, 306–317.

(10) Berendsen, H. J. C.; van der Spoel, D.; van Drunen, R. GROMACS: A Message-Passing Parallel Molecular Dynamics Implementation. *Comput. Phys. Commun.* **1995**, *91*, 43–56.

(11) Schmid, N.; Eichenberger, A. P.; Choutko, A.; Riniker, S.; Winger, M.; Mark, A. E.; Gunsteren, W. F. Definition and Testing of the GROMOS Force-Field Versions 54A7 and 54B7. *Eur. Biophys. J.* **2011**, *40*, 843–856.

(12) Ferreira, T. M.; Coreta-Gomes, F.; Ollila, O. H. S.; Moreno, M. J.; Vaz, W. L. C.; Topgaard, D. Cholesterol and POPC Segmental Order Parameters in Lipid Membranes: Solid State 1 H– 13 C NMR and MD Simulation Studies. *Phys. Chem. Chem. Phys.* **2013**, *15* (6), 1976–1989. https://doi.org/10.1039/C2CP42738A.

(13) Ferreira, T. M.; Sood, R.; Bärenwald, R.; Carlström, G.; Topgaard, D.; Saalwächter, K.; Kinnunen, P. K. J.; Ollila, O. H. S. Acyl Chain Disorder and Azelaoyl Orientation in Lipid Membranes Containing Oxidized Lipids. *Langmuir* **2016**, *32* (25), 6524–6533. https://doi.org/10.1021/acs.langmuir.6b00788.

(14) Berger, O.; Edholm, O.; Jähnig, F. Molecular Dynamics Simulations of a Fluid Bilayer of Dipalmitoylphosphatidylcholine at Full Hydration, Constant Pressure, and Constant Temperature. *Biophys. J.* **1997**, *72* (5), 2002–2013. https://doi.org/10.1016/S0006-3495(97)78845-3.

(15) Bachar, M.; Brunelle, P.; Tieleman, D. P.; Rauk, A. Molecular Dynamics Simulation of a Polyunsaturated Lipid Bilayer Susceptible to Lipid Peroxidation. *J. Phys. Chem. B* **2010**, *114* (50), 17002–17002. https://doi.org/10.1021/jp110570m.

(16) Ollila, S.; Hyvönen, M. T.; Vattulainen, I. Polyunsaturation in Lipid Membranes: Dynamic Properties and Lateral Pressure Profiles. *J. Phys. Chem. B* **2007**, *111* (12), 3139–3150. https://doi.org/10.1021/jp065424f.

(17) Höltje, M.; Förster, T.; Brandt, B.; Engels, T.; Von Rybinski, W.; Höltje, H. D. Molecular Dynamics Simulations of Stratum Corneum Lipid Models: Fatty Acids and Cholesterol. *Biochim. Biophys. Acta - Biomembr.* **2001**, *1511* (1), 156–167. https://doi.org/10.1016/S0005-2736(01)00270-X.

(18) Tieleman, D. P.; MacCallum, J. L.; Ash, W. L.; Kandt, C.; Xu, Z.; Monticelli, L. Membrane Protein Simulations with a United-Atom Lipid and All-Atom Protein Model: Lipid-Protein Interactions, Side Chain Transfer Free Energies and Model Proteins. *J. Phys. Condens. Matter* **2006**, *18* (28). https://doi.org/10.1088/0953-8984/18/28/S07.

(19) Khandelia, H.; Mouritsen, O. G. Lipid Gymnastics: Evidence of Complete Acyl Chain

Reversal in Oxidized Phospholipids from Molecular Simulations. *Biophys. J.* **2009**, *96* (7), 2734–2743. https://doi.org/10.1016/j.bpj.2009.01.007.

(20) Zhao, W.; Róg, T.; Gurtovenko, A. A.; Vattulainen, I.; Karttunen, M. Atomic-Scale Structure and Electrostatics of Anionic Palmitoyloleoylphosphatidylglycerol Lipid Bilayers with Na+ Counterions. *Biophys. J.* **2007**, *92* (4), 1114–1124. https://doi.org/10.1529/biophysj.106.086272.

(21) Ollila, O. H. S.; Ferreira, T. M.; Topgaard, D. MD simulation trajectory and related files for POPC bilayer (Berger model delivered by Tieleman, Gromacs 4.5) https://zenodo.org/record/13279 (accessed 2019 -09 -12). https://doi.org/10.5281/ZENODO.13279.

(22) Ollila, O. H. S.; Ferreira, T.; Topgaard, D. MD simulation trajectory and related files for POPC/cholesterol (15 mol%) bilayer (Berger model delivered by Tieleman, modified Höltje, Gromacs 4.5) https://zenodo.org/record/13281 (accessed 2019 -09 -12). https://doi.org/10.5281/zenodo.13281.

(23) Ollila, O. H. S. MD simulation trajectory and related files for POPC bilayer with 30 mol% of protonated pazePC (Berger, Gromacs 4.5.) https://zenodo.org/record/44660 (accessed 2019 -09 -12). https://doi.org/10.5281/zenodo.44660.

(24) Ollila, Samuli; Khandelia, H. MD simulation trajectory and related files for POPC bilayer with 30 mol% of deprotonated pazePC (Berger, Gromacs 4.5.) https://zenodo.org/record/44622 (accessed 2019 -09 -12). https://doi.org/10.5281/zenodo.44622.

(25) Ghahremanpour, M. M.; Arab, S. S.; Aghazadeh, S. B.; Zhang, J.; Van Der Spoel, D. MemBuilder: A Web-Based Graphical Interface to Build Heterogeneously Mixed Membrane Bilayers for the GROMACS Biomolecular Simulation Program. *Bioinformatics* **2014**, *30* (3), 439–441. https://doi.org/10.1093/bioinformatics/btt680.

(26) Delaglio, F. PDB Utility Server. Create an Extended Structure https://spin.niddk.nih.gov/bax/nmrserver/pdbutil/ext.html.

(27) Crescenzi, O.; Tomaselli, S.; Guerrini, R.; Salvadori, S.; D'Ursi, A. M.; Temussi, P. A.; Picone, D. Solution Structure of the Alzheimer Amyloid β-Peptide (1-42) in an Apolar Microenvironment: Similarity with a Virus Fusion Domain. *Eur. J. Biochem.* **2002**, *269* (22),

5642–5648. https://doi.org/10.1046/j.1432-1033.2002.03271.x.

(28) Prasad, A. K.; Tiwari, C.; Ray, S.; Holden, S.; Armstrong, D. A.; Rosengren, K. J.; Rodger, A.; Panwar, A. S.; Martin, L. L. Secondary Structure Transitions for a Family of Amyloidogenic, Antimicrobial Uperin 3 Peptides in Contact with Sodium Dodecyl Sulfate. *Chempluschem* **2022**, *87* (1), e202100408. https://doi.org/10.1002/cplu.202100408.

(29) Vivekanandan, S.; Brender, J. R.; Lee, S. Y.; Ramamoorthy, A. A Partially Folded Structure of Amyloid-Beta(1-40) in an Aqueous Environment. *Biochem. Biophys. Res. Commun.* **2011**, *411* (2), 312–316. https://doi.org/10.1016/j.bbrc.2011.06.133.

(30) Essmann, U.; Perera, L.; Berkowitz, M. L.; Darden, T.; Lee, H.; Pedersen, L. G. A Smooth Particle Mesh Ewald Method. *J. Chem. Phys.* **1995**, *103*, 8577–8593.

(31) Darden, T.; York, D.; Pedersen, L. Particle Mesh Ewald: An N ·log( N ) Method for Ewald Sums in Large Systems. *J. Chem. Phys.* **1993**, *98* (12), 10089–10092. https://doi.org/10.1063/1.464397.

(32) Hess, B. P-LINCS: A Parallel Linear Constraint Solver for Molecular Simulation. *J. Chem. Theory Comput.* **2008**, *4* (1), 116–122. https://doi.org/10.1021/ct700200b.

(33) Berendsen, H. J. C.; Postma, J. P. M.; van Gunsteren, W. F.; Hermans, J. Interaction Models for Water in Relation to Protein Hydration. In *The Jerusalem Symposia on Quantum Chemistry and Biochemistry*; Pullman, B., Ed.; Reidel: Dordrecht, 1981; pp 331–342.

(34) Miyamoto, S.; Kollman, P. A. SETTLE: An Analytical Version of the SHAKE and RATTLE Algorithm for Rigid Water Models. *J. Comput. Chem.* **1992**, *13* (8), 952–962. https://doi.org/10.1002/jcc.540130805.

(35) Bussi, G.; Donadio, D.; Parrinello, M. Canonical Sampling through Velocity Rescaling. *J. Chem. Phys.* **2007**, *126*, 14101.

(36) Berendsen, H. J. C.; Postma, J. P. M.; van Gunsteren, W. F.; DiNola, A.; Haak, J. R. Molecular Dynamics with Coupling to an External Bath. *J. Chem. Phys.* **1984**, *81*, 3684–3690.

(37) Humphrey, W.; Dalke, A.; Schulten, K. VMD: Visual Molecular Dynamics. *J. Mol. Graph.* **1996**, *14* (1), 33–38. https://doi.org/10.1016/0263-7855(96)00018-5.

(38) Daura, X.; Gademann, K.; Jaun, B.; Seebach, D.; van Gunsteren, W. F.; Mark, A. E. Peptide Folding: When Simulation Meets Experiment. *Angew. Chemie Int. Ed.* **1999**, *38* (1–2), 236–

240. https://doi.org/10.1002/(SICI)1521-3773(19990115)38:1/2<236::AID-ANIE236>3.0.CO;2-M.

(39) Kabsch, W.; Sander, C. Dictionary of Protein Secondary Structure - Pattern-Recognition of Hydrogen-Bonded and Geometrical Features. *Biopolymers* **1983**, *22* (12), 2577–2637. https://doi.org/10.1002/bip.360221211.

(40) Touw, W. G.; Baakman, C.; Black, J.; Te Beek, T. A. H.; Krieger, E.; Joosten, R. P.; Vriend, G. A Series of PDB-Related Databanks for Everyday Needs. *Nucleic Acids Res.* **2015**, *43* (D1), D364–D368. https://doi.org/10.1093/nar/gku1028.

(41) Mattila, J. P.; Sabatini, K.; Kinnunen, P. K. J. Oxidized Phospholipids as Potential Novel Drug Targets. *Biophys. J.* **2007**, *93* (9), 3105–3112. https://doi.org/10.1529/biophysj.107.103887.

(42) Pande, A. H.; Kar, S.; Tripathy, R. K. Oxidatively Modified Fatty Acyl Chain Determines Physicochemical Properties of Aggregates of Oxidized Phospholipids. *Biochim. Biophys. Acta - Biomembr.* **2010**, *1798* (3), 442–452. https://doi.org/10.1016/j.bbamem.2009.12.028.

(43) Haberland, M. E.; Reynolds, J. A. Self Association of Cholesterol in Aqueous Solution. *Proc. Natl. Acad. Sci. U. S. A.* **1973**, *70* (8), 2313–2316. https://doi.org/10.1073/pnas.70.8.2313.

(44) Avanti. Critical Micelle Concentrations (CMCs) https://avantilipids.com/tech-support/physical-properties/cmcs.
